# Transcriptional response of *Candida auris* to the Mrr1 inducers methylglyoxal and benomyl

**DOI:** 10.1101/2022.03.02.482751

**Authors:** Amy R. Biermann, Deborah A. Hogan

**Author notes:** To whom correspondence should be addressed Department of Microbiology and Immunology, Geisel School of Medicine at Dartmouth Rm 208 Vail Building, Hanover, NH 03755.

## Abstract

*Candida auris* is an urgent threat to human health due to its rapid spread in healthcare settings and its repeated development of multidrug resistance. Diseases that put individuals at a higher risk for *C. auris* infection, such as diabetes, kidney failure, or immunocompromising conditions, are associated with elevated levels of methylglyoxal (MG), a reactive dicarbonyl compound derived from several metabolic processes. In other *Candida* species, expression of MG reductase enzymes that catabolize and detoxify MG are controlled by Mrr1, a multidrug resistance-associated transcription factor, and MG induces Mrr1 activity. Here, we used transcriptomics and genetic assays to determine that *C. auris MRR1a* contributes to MG resistance, and that the main Mrr1a targets are an MG reductase and *MDR1*, which encodes an drug efflux protein. The *C. auris* Mrr1a regulon is smaller than Mrr1 regulons described in other species. In addition to MG, benomyl (BEN), a known Mrr1 stimulus, induces *C. auris* Mrr1 activity, and characterization of the *MRR1a*-dependent and independent transcriptional responses revealed substantial overlap in genes that were differentially expressed in response to each compound. Additionally, we found that an *MRR1* allele specific to one *C. auris* phylogenetic clade, clade III, encodes a hyperactive Mrr1 variant, and this activity correlated with higher MG resistance. *C. auris MRR1a* alleles were functional in *Candida lusitaniae* and were inducible by BEN, but not by MG, suggesting that the two Mrr1 inducers act via different mechanisms. Together, the data presented in this work contribute to the understanding Mrr1 activity and MG resistance in *C. auris*.

**Importance:** *Candida auris* is a fungal pathogen that has spread since its identification in 2009 and is of concern due to its high incidence of resistance against multiple classes of antifungal drugs. In other *Candida* species, the transcription factor Mrr1 plays a major role in resistance against azole antifungals and other toxins. More recently, Mrr1 has been recognized to contribute to resistance to methylglyoxal (MG), a toxic metabolic byproduct. Here, we show that *C. auris MRR1a*, the closest ortholog to *MRR1* in other species, contributes to resistance to MG, and that Mrr1a strongly co-regulates expression of *MGD1*, encoding a methylglyoxal reductase enzyme and *MDR1*, encoding an efflux protein involved in resistance to azole drugs, antimicrobial peptides and bacterial products. We found that one major clade of *C. auris* has a constitutively active Mrr1 despite high azole resistance due to other mutations, and that this high Mrr1a activity correlates with higher MG resistance. Finally, we gain insights into the activities of MG and another Mrr1 inducer, benomyl, to better understand *C. auris* regulation of phenotypes relevant *in vivo*.

## Introduction

Although *Candida albicans* has historically been the most prominent *Candida* species associated with both superficial and invasive fungal infections, worldwide incidence of non-albicans *Candida* (NAC) species is increasing (1–10). Of particular concern is *Candida auris*, which the CDC classifies as an urgent threat due to its relatively high frequency of resistance to multiple different classes of drugs including amphotericin B, echinocandins and azoles (reviewed in (11)). Since its recognition as a novel *Candida* species in 2009, *C. auris*, has been reported in at least 40 countries (12–14). Whole-genome sequencing (WGS) analyses of *C. auris* isolates collected from across the globe indicate the concurrent emergence of four genetically distinct clades (15) with a potential fifth clade defined more recently (16). *C. auris* is thought to primarily colonize the skin (17–19) in addition to a diverse array of body sites, and most clinical isolates to date have been isolated from blood (20). Once *C. auris* has disseminated to the bloodstream, it can cause potentially fatal candidemia which has an estimated global mortality rate ranging from about 30 – 60% (15, 21, 22).

The resistance to azoles in *C. auris* is multifactorial; it has been shown that certain mutations in *ERG11* (15, 23–31) and overproduction of Cdr1p (32–36) contribute to resistance to fluconazole (FLZ). In multiple *Candida* species, the transcriptional regulator Mrr1 also plays a role in FLZ resistance (37–45), and Mayr and colleagues (46) found three *C. auris* homologs of the transcriptional regulator Mrr1, and showed that one of them, *MRR1a*, modestly affected fluconazole resistance. Previously, we demonstrated that in *Candida* (*Clavispora*) *lusitaniae*, which is more closely related to *C. auris* relative to other well-studied *Candida* species (12, 47), Mrr1 regulates the expression of *MDR1*, and overexpression of *MDR1* confers resistance to FLZ (40, 48–55), the host antimicrobial peptide histatin-5 (40, 56), bacterially produced phenazines (40), and other toxic compounds (57) in multiple *Candida* species. *C. lusitaniae* Mrr1 also regulates dozens of other genes with two of the most strongly regulated genes encoding methylglyoxal (MG) reductase enzymes, *MGD1* and *MGD2* (37, 40, 58). Mrr1 contributes to *C. lusitaniae* resistance to MG (58), which is a spontaneously formed dicarbonyl electrophile generated as a byproduct of several metabolic processes by all living cells (reviewed in (59)). Via its carbonyl groups, MG reacts non-enzymatically with biomolecules, which can lead to cellular stress and toxicity (reviewed in (59)). Some of the risk factors (60–69) for candidiasis caused by *C. auris* or other *Candida* spp., such as diabetes (70–72), kidney disease (73–76), or septic shock (77), are associated with elevated MG in human serum. MG resistance across clinical isolates of the same *Candida* species, including *C. auris*, can vary (58).

Through specific regulators, MG and other reactive electrophiles induce stress responses in bacteria (78–80), plants (reviewed in (81)), mammals (reviewed in (82)), and the yeasts *Saccharomyces cerevisiae* (83–87) and *Schizosaccharomyces pombe* (88, 89) at subinhibitory concentrations. We found in *C. lusitaniae*, MG induces expression of *MGD1* and *MGD2* as well as *MDR1*, through a mechanism that involved Mrr1 (58), and that MG increased fluconazole (FLZ) resistance. *C. auris* displays nosocomial transmission (61-63, 65-69), in part due to its resistance to high temperatures (90) and common surface antiseptics (91), and persistence on abiotic surfaces including latex and nitrile gloves (92), plastics (90), and axillary temperature probes (93). The factors that control *C. auris* stress resistance are not yet known.

In the present study, we show that *C. auris MRR1a* regulates resistance to MG and that MG is an inducer of Mrr1-regulated gene expression. Mrr1a regulates the gene orthologous to the methylglyoxal reductase genes *C. lusitanaie MGD1* in addition to *MDR1*, which regulates FLZ efflux, but the Mrr1a regulon is smaller than that described for other species. Furthermore, we characterize Mrr1a in both Clade I and Clade III isolates and show that the Mrr1 variant in Clade III is constitutively active. Transcriptomics analysis shows that MG elicits a large transcriptional response that is similar in both Clade I and Clade III, and that there are commonalities in the responses elicited by MG and the Mrr1 inducer benomyl. These data support the model that Mrr1 is a regulator of MG resistance in coordination with efflux proteins such as Mdr1 and provides the basis for future studies on the roles of Mrr1 and MG in survival of *C. auris* in hospital settings.

## Results

### Mrr1a regulates expression of orthologs to *MDR1* and *MGD1* in *C. auris* strain B11221 and is involved in MG resistance

To determine whether the *C. auris MRR1* orthologs *MRR1a*, *MRR1b*, and *MRR1c* contributed to resistance to MG, we performed growth kinetic assays in YPD +/- 5 mM, 10 mM, or 15 mM MG. At MG concentrations of 10 mM (Fig. 1A) and 15 mM (Fig. S1), the *mrr1a*Δ mutant displayed a substantial growth defect relative to WT, while the *mrr1b*Δ and *mrr1c*Δ mutants exhibited growth comparable to WT. None of the mutants (*mrr1a*Δ, *mrr1b*Δ, and *mrr1c*Δ) differed from the parental isolate B11221 (WT) in YPD alone or in the presence of 5 mM MG (Fig. S1). Like *C. lusitaniae*, the *C. auris* genome encodes multiple putative MG reductases; the closest orthologs to *MGD1* and *MGD2* were *CJI97_000658* and *CJI97_004624*, respectively, in the B11221 genome assembly (58) and we will henceforth refer to these genes as *MGD1* and *MGD2*. For reference, *MGD1* and *MGD2* correspond to B9J08_000656 and B9J08_004828 respectively. By quantitative real-time PCR (qRT-PCR), basal expression of *MGD1* was significantly decreased 24-fold in the *mrr1a*Δ mutant relative to B11221 WT (Fig. 1B), and expression of *MGD2* trended lower in the *mrr1a*Δ mutant (∼1.2-fold) but this difference did not reach statistical significance (Fig. 1C). *MGD1* was also more highly expressed than *MGD2* in the WT B11221 as in *C. lusitaniae* (58). Consistent with the transcriptional patterns, *C. auris* Mgd1 shares slightly more identity with *C. lusitaniae* Mgd1 than does *C. auris* Mgd2 (63% identity versus 61% identity).

**Figure 1.**
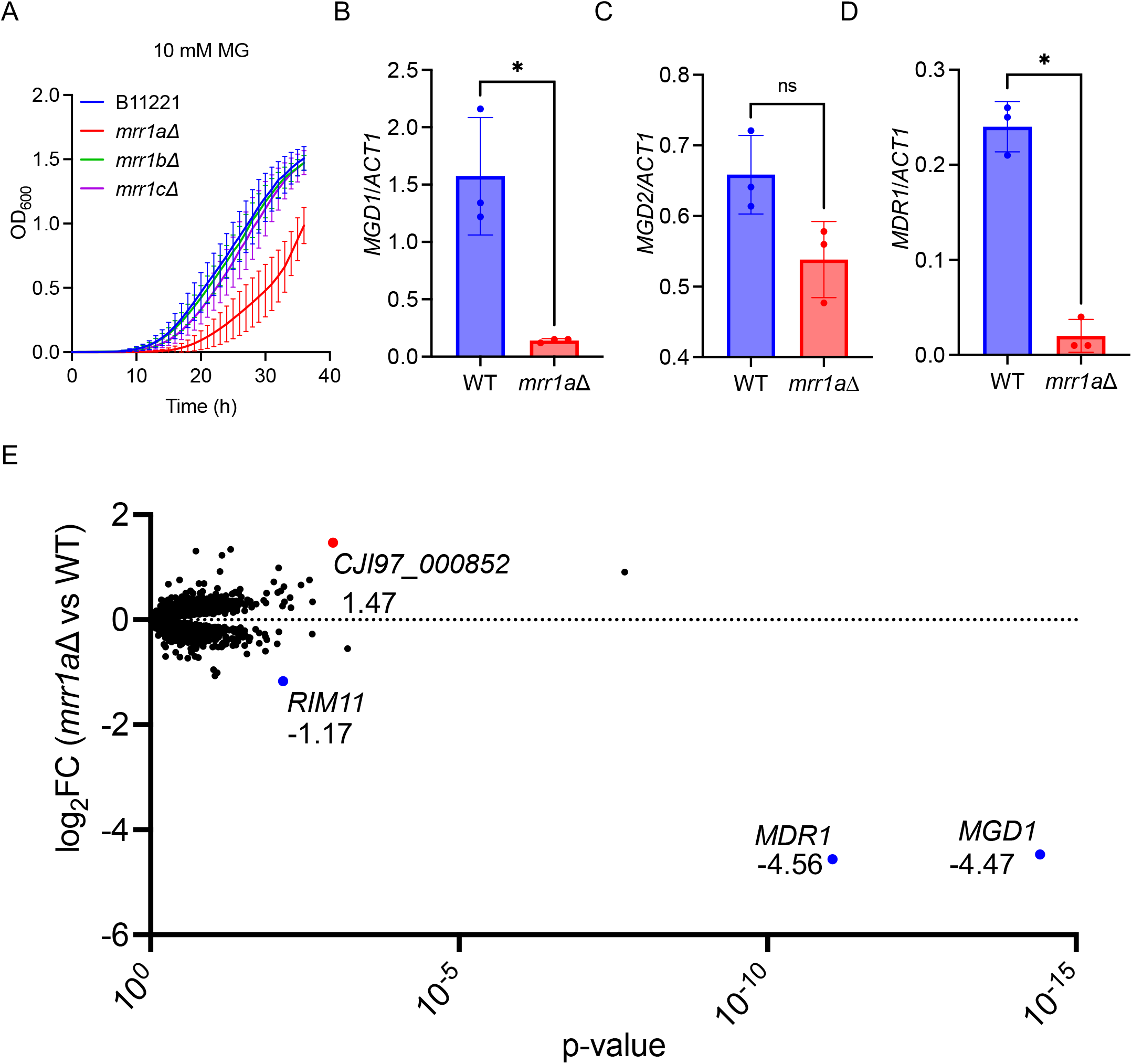
Mrr1a regulates expression of *MGD1* and *MDR1* in *C. auris* isolate B11221. **A)** Growth curves of B11221 WT (blue) and its *mrr1a*Δ (red), *mrr1b*Δ (green), and *mrr1c*Δ (purple) derivatives in YPD + 10 mM MG. Data shown represent the mean ± SD for three independent experiments. **B-C)** qRT-PCR assessment of *MGD1* **(B)** and *MDR1* **(C)** expression in B11221 WT (blue) and *mrr1a*Δ (red) cultures grown to exponential phase in YPD at 37°C. Data shown represent the mean ± SD for three independent experiments. Ratio paired t-test was used for statistical evaluation; * p < 0.05. **D)** Volcano plot of all quantified genes in B11221 WT vs *mrr1a*Δ in the control condition. Each point represents a single gene; blue points indicate genes significantly more highly expressed in WT; red points indicate genes significantly more highly expressed in *mrr1a*Δ. Numbers adjacent to each colored point indicate the log_2_FC in *mrr1a*Δ versus WT.

In the *C. auris* B11221 background, expression of *MDR1*, another target of Mrr1 in other species including *C. lusitaniae*, also depended on Mrr1a, as the *mrr1a*Δ mutant exhibited a significant 21-fold decrease in *MDR1* expression compared to the WT parent (Fig. 1D). These results indicate that in *C. auris MDR1* and *MGD1* are co-regulated, as has been reported in *C. albicans* (44, 45, 94–96), *C. parapsilosis* (97), and *C. lusitaniae* (37, 39, 40, 58, 98) and that higher expression of *MGD1* and/or *MDR1* contributes to growth in high concentrations of MG (Fig. 1A).

In *C. lusitaniae* and other *Candida* species, Mrr1 regulates dozens of genes in addition to *MDR1* and *MGD1* (37, 40). To further elucidate the Mrr1a regulon in *C. auris* isolate B11221, we performed an RNA-seq analysis of in B11221 WT and its *mrr1a*Δ derivative in cells from exponential phase cultures grown at 37°C in YPD. In the control condition (YPD + dH_2_O), only four genes, including *MDR1* and *MGD1*, were differentially expressed between the two strains with the cutoff of a log_2_ fold change (log_2_FC) ≥ 1.00 or ≤ −1.00 and a p-value less than 0.05 (Fig 1E and **Supplementary File S1** for all data). *MGD1* and *MDR1* showed a 22- and 24-fold decrease, respectively, in *mrr1a*Δ compared to WT, consistent with our qRT-PCR data. *CJI97_005632*, which was 2.25-fold lower in *mrr1*aΔ, is orthologous to the *C. albicans* genes *RIM11* and *C2_04280W_A*, both of which are predicted to encode proteins with serine/threonine kinase activity, though it is worth noting that levels of the transcript were much lower than levels of *MDR1* and *MGD1*. *CJI97_000852*, which was 2.77-fold higher in *mrr1a*Δ than in WT, has 16 orthologs of diverse predicted or known functions in *C. albicans*, including *USO5*, *USO6*, and *RBF1* (Fig. 1E and **Supplementary File S1**). Notably, *MGD2* was not differentially expressed between B11221 WT and the *mrr1a*Δ mutant in our RNA-seq data (**Supplementary File S1**), consistent with our qRT-PCR results described above.

### Mrr1a regulates only *MDR1* and *MGD1* in response to MG and benomyl

We have previously shown in *C. lusitaniae* that MG induces expression of the Mrr1-regulated genes *MGD1* and *MGD2* in an Mrr1-dependent manner, and *MDR1* in a partially Mrr1-dependent manner (58). To determine if MG would induce expression of *MGD1*, *MGD2*, and/or *MDR1* in *C. auris*, we purified RNA for qRT-PCR from exponential-phase cultures of B11221 WT and *mrr1a*Δ treated with 5 mM MG or an equal volume of dH_2_O for 15 minutes. We found that MG treatment significantly enhanced expression of *MGD1* in WT by 2.4-fold but not in *mrr1a*Δ (Fig. 2A). *MGD1* was also induced by a 30-min treatment with 25 µg/mL benomyl (BEN), a known inducer of Mrr1-regulated genes in other *Candida* species (37, 41, 43, 95, 99–104), by 7.5-fold in the WT (Fig. 2A). The different treatment times for MG and BEN were used to be consistent with previous studies using either compound in the related species *C. lusitaniae* (37, 58). Expression of *MDR1* was also more highly induced by treatment with either MG or BEN in WT compared to the *mrr1a*Δ mutant by 6- and 14.5-fold respectively (Fig. 2B). Although *MDR1* expression was significantly induced by MG and BEN in the *mrr1a*Δ, transcript levels of *MDR1* were approximately 20-fold higher in the WT than in the *mrr1a*Δ under these conditions (Fig. 2B), suggesting that Mrr1a is required for maximum expression of *MDR1* in response to stimuli.

**Figure 2.**
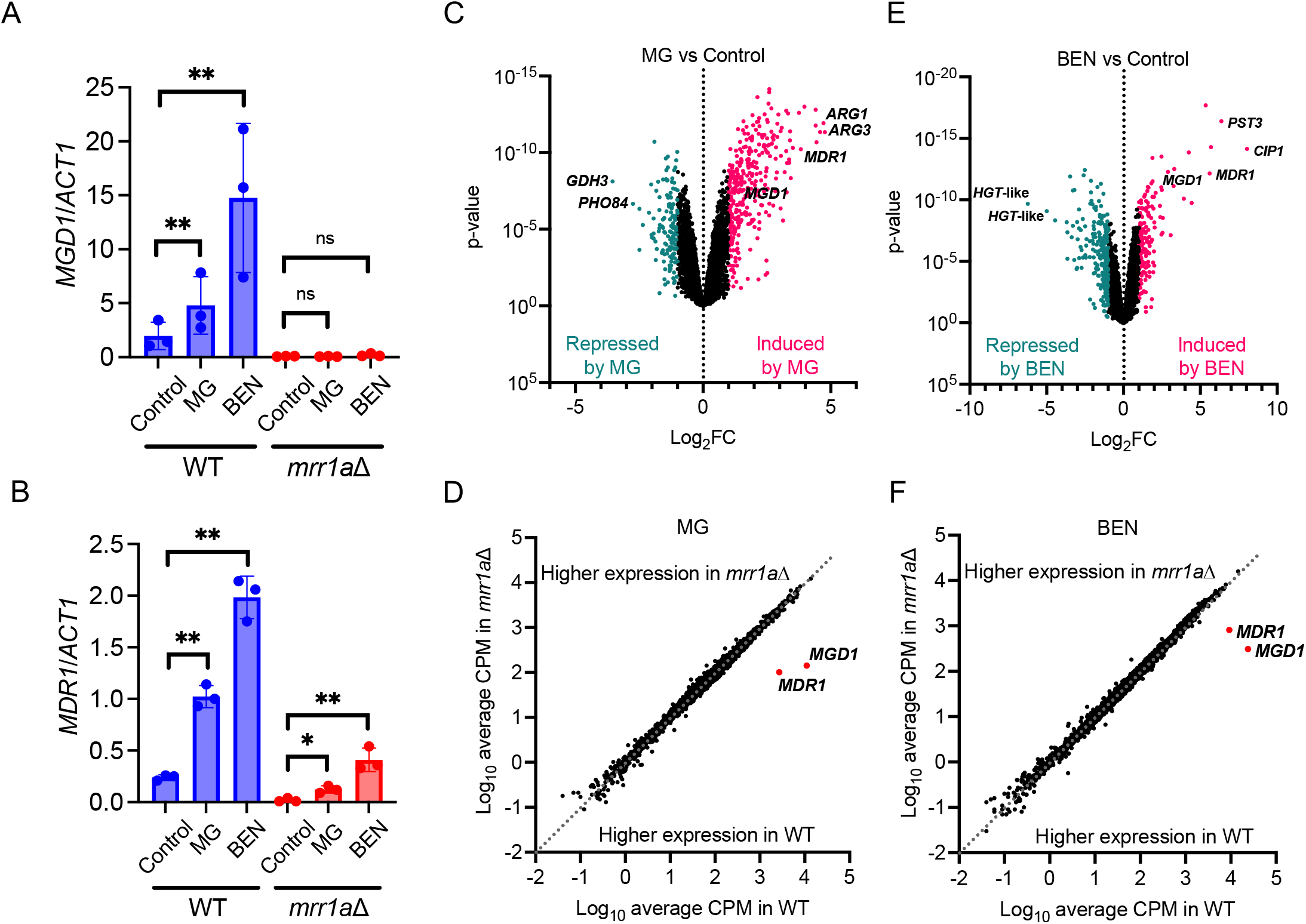
MG and BEN both lead to a vast transcriptional response in *C. auris* B11221, which includes upregulation of *MDR1* and *MGD1*. **A-B)** qRT-PCR analysis for expression of *MGD1* **(A)** and *MDR1* **(B)** in exponential-phase cultures of B11221 WT (blue) or *mrr1a*Δ (red) treated with MG or BEN as indicated. Data shown represent the mean ± SD for three independent experiments. Ratio paired t-test was used for statistical evaluation; ns p > 0.05, * p < 0.05, ** p < 0.01. **C-D)** Volcano plots of all quantified genes in B11221 WT treated with either MG **(C)** or BEN **(D)**. Each point represents a single gene; magenta points indicate genes that were significantly upregulated compared to the control condition, teal points indicate genes that were significantly downregulated compared to the control condition. *MDR1* and *MGD1* are shown along with the two most up- and down-regulated genes in each condition. **E-F)** Scatter plots of the average CPMs of all quantified genes in *mrr1a*Δ vs. B11221 WT treated with MG **(E)** or BEN **(F)**. Each point represents a single gene. Points below the dotted line indicate genes that were more highly expressed in the WT, and points above the dotted line indicated genes that were more highly expressed in the *mrr1a*Δ mutant. *MDR1* and *MGD1* are shown with red dots for reference.

To describe the complete Mrr1-dependent MG- and BEN-response regulon under our test conditions in *C. auris*, we also performed RNA-seq on exponential-phase cultures of B11221 WT and *mrr1a*Δ treated with MG or BEN as described above. In B11221 WT, MG led to the upregulation of 319 genes and downregulation of 133 genes compared to the control condition (Fig. 2C and **Supplementary File S2**). In the *mrr1a*Δ mutant, MG led to the upregulation of 349 genes and downregulation of 143 genes compared to the control condition (Fig. S2A and **Supplementary File S2**). Consistent with our qRT-PCR data in Fig. 2A, MG induced expression of *MGD1* in the WT but not in the *mrr1a*Δ mutant (Table S1 and **Supplementary File S2**). Although expression of *MDR1* was significantly induced by MG in both the WT and the *mrr1a*Δ mutant (Table S1 and **Supplementary File S2**), levels of *MDR1* were substantially lower in the *mrr1a*Δ mutant even in the presence of MG (Fig. 2D and **Supplementary File S2**), also in agreement with our qRT-PCR data. *MGD1* and *MDR1* strongly stood out as the only two genes in the MG response that were strongly dependent on Mrr1a (Fig. 2D).

Treatment with BEN led to upregulation of 160 genes and downregulation of 163 genes in the WT (Fig. 2E and **Supplementary File S3**). In the *mrr1a*Δ mutant, 181 genes were upregulated, and 229 genes were downregulated in response to BEN (Fig. S2B and **Supplementary File S3**). Like MG, induction of *MGD1* by BEN was completely dependent on Mrr1a (Table S1 and **Supplementary File S3**) and *MGD2* expression was not induced by BEN (**Supplementary File S3**). Expression of *MDR1* was also induced by BEN in both the WT and the *mrr1a*Δ mutant, but as with MG, *MDR1* levels in the *mrr1a*Δ mutant did not reach that of the WT even with BEN treatment (Fig. 2F and **Supplementary File S3**). Again, *MGD1* and *MDR1*, appear to be the only genes in *C. auris* whose induction of expression by either MG or BEN is dependent on Mrr1a. The Mrr1a-independent responses to MG and BEN are discussed further below.

### B11221 has higher basal expression of *MDR1* and of putative MG reductase genes compared to the Clade I isolate AR0390

Many Clade III isolates, including B11221, contain an N647T single nucleotide polymorphism (SNP) in *MRR1a* (25, 105). In (105), this SNP was proposed to be a gain-of-function mutation due to the resistance of Clade III isolates against azoffluxin, a novel antifungal compound that inhibits expression and activity of *C. auris* efflux pumps. As a first step to determine if there were activity differences between the Mrr1a variant that was found Clade III strains was different from that encoded by the alleles found in Clade I, II, and IV strains, we compared MG sensitivity of B11221 to that of Clade I isolate AR0390. Interestingly, AR0390 grew substantially better than B11221 in the YPD control but showed a greater reduction in growth in YPD with 5 mM MG than did B11221 (Fig. S3). At concentrations of 10 mM (Fig. 3A) and 15 mM MG (Fig. S3), AR0390 exhibited a profound growth defect compared to B11221. To determine if differences in MG sensitivity were due to differences in *MGD1* expression, we measured basal expression of *MGD1* and its co-regulated gene *MDR1* in B11221 and AR0390 using qRT-PCR. Both genes were significantly more highly expressed in B11221 by 42- and 4.2-fold, respectively (Fig. 3B-C).

**Figure 3.**
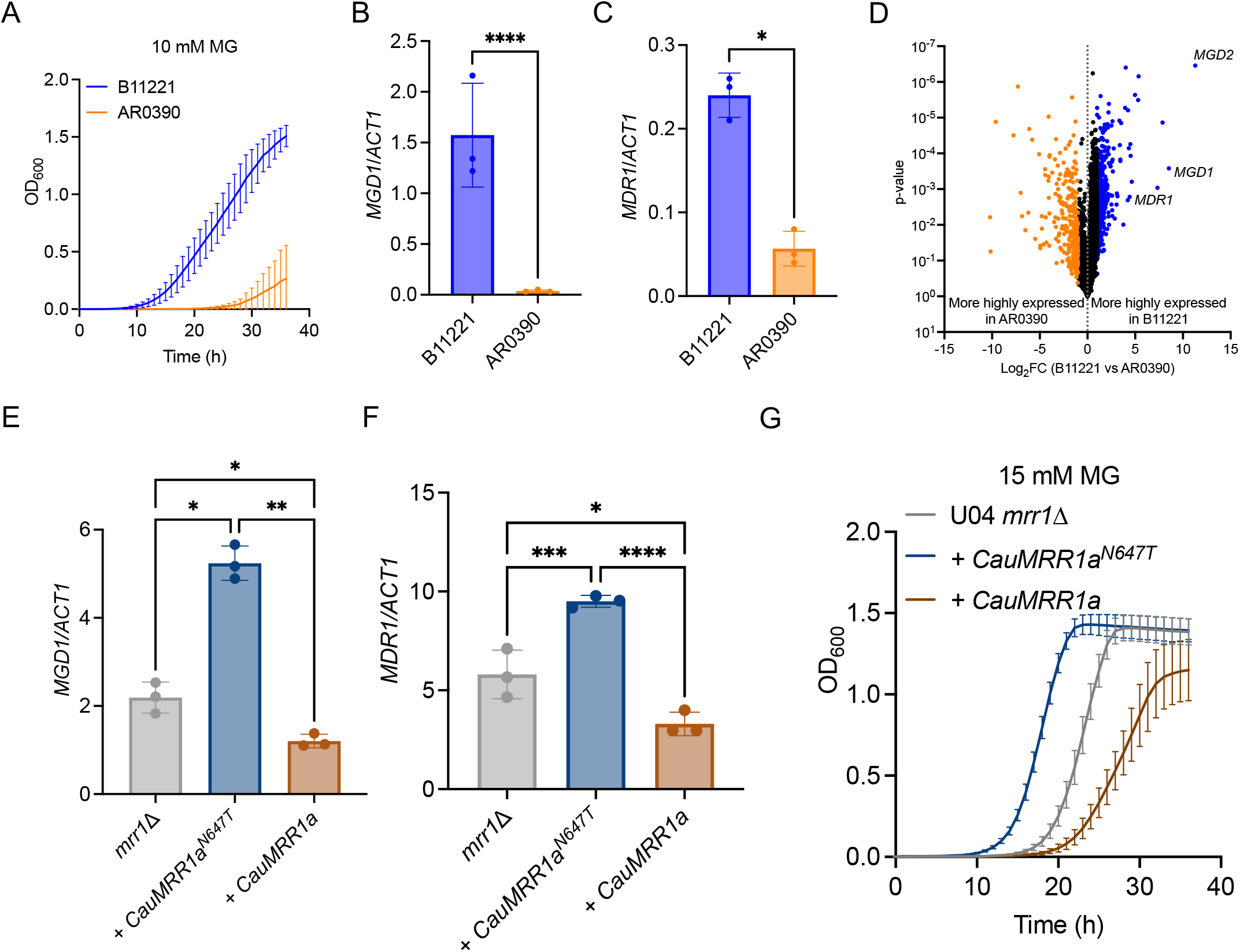
*MDR1* and *MGD1* are among the genes significantly more highly expressed in isolate B11221 compared to isolate AR0390. **A)** Growth curves of B11221 (blue) and AR#0390 (orange) in YPD + 10 mM MG. Data shown represent the mean ± SD for three independent experiments. **B-C)** qRT-PCR assessment of *MGD1* **(B)** and *MDR1* **(C)** expression in B11221 (blue) and AR0390 (orange) grown to exponential phase in YPD at 37°C. Data shown represent the mean ± SD for three independent experiments. Ratio paired t-test was used for statistical evaluation; * p < 0.05, **** p < 0.0001. **D)** Volcano plot of all quantified genes, matched by syntenic ortholog, in B11221 and AR0390 in the control condition (YPD). Each point represents a single gene; blue points indicate genes significantly more highly expressed in B11221; orange points indicate genes significantly more highly expressed in AR0390. **E-F)** qRT-PCR expression analysis for *MGD1* **(E)** and *MDR1* **(F)** in *C. lusitaniae* U04 *mrr1*Δ (grey) and its derivatives expressing *CauMRR1a^N647T^* (dark blue) or *CauMRR1a* (brown). Data shown represent the mean ± SD for three independent experiments. One-way ANOVA was used for statistical evaluation; * p < 0.05, ** p < 0.01, *** p < 0.001, **** p < 0.0001. **G)** Growth curves of *C. lusitaniae* U04 *mrr1*Δ (grey) and its derivatives expressing *CauMRR1a^N647T^* (dark blue) or *CauMRR1a* (brown) in YPD + 15 mM MG. One representative experiment of three independent experiments is shown; error bars represent the standard deviation of technical replicates within the experiment.

To gain a deeper understanding of the broader transcriptional differences between B11221 and AR0390, we compared the basal global gene expression in YPD of the two strains using RNA-seq. First, we matched the 5227 syntenic orthologs between the genomes of B11221 and the Clade I reference strain B8441 to compare expression of each gene under the control condition. Of these, 755 genes were differentially expressed between B11221 and AR0390 in the control condition (|log_2_FC| ≥ 1.00, FDR-corrected p < 0.05) (Fig. 3D**, Supplementary File S4)**. The top twenty differentially expressed genes whose orthologs have known or predicted functions in *C. albicans* are reported in Table S3. Strikingly, the two genes which exhibited the largest difference in expression between B11221 and AR0390 were *MGD2* (log_2_FC = 11.29) and *MGD1* (log_2_FC = 8.53) (Fig. 3D, Table S3, and **Supplementary File S4**). A third gene with homology to MG reductases, *CJI97_001800*/*B9J08_002257*, was also more highly expressed in B11221, although the log_2_FC in expression of this gene in B11221 vs AR0390 was only 1.41 (**Supplementary File S4**). Low expression of *MGD1*, *MGD2*, and/or *B9J08_002257* may contribute to the severe growth defect of AR0390 in the presence of MG. Consistent with our qRT-PCR data, *MDR1* was also significantly more highly expressed in B11221 relative to AR0390 (log_2_FC = 4.42) (Fig. 3D and Table S3). Although *MGD2* and *B9J08_002257* do not appear to be regulated by Mrr1a in our studies, it is nonetheless interesting to note the elevated expression of three putative MG reductases in the *MDR1*-overexpressing *C. auris* isolate B11221, as the co-expression of *MDR1* with at least one MG reductase has been reported in numerous studies in other *Candida* species (37, 40, 44, 45, 58, 94–97).

### Clade III Mrr1a^N647T^ exhibits a gain-of-function phenotype compared to Clade I Mrr1a when expressed in *C. lusitaniae*

To compare the activities of the proteins encoded by the *MRR1a* alleles of B11221 and AR0390 more directly, we heterologously expressed each allele, henceforth referred to as *CauMRR1a^N647T^* and *CauMRR1a* respectively, independently in a *C. lusitaniae mrr1*Δ mutant previously generated and characterized by our lab (37, 40, 58). All three *C. lusitaniae* clones expressing *CauMRR1a^N647T^* which we tested exhibited a four-fold increase in fluconazole (FLZ) MIC relative to the U04 *mrr1*Δ parent (16 µg/mL versus 4 µg/mL, Table S4), confirming that *C. auris* Clade III *MRR1a* can complement *MRR1*-dependent FLZ resistance in *C. lusitaniae* and adding support to the hypothesis that the N647T substitution in Clade III *MRR1a* confers increased activity. However, the FLZ MIC of the three tested *C. lusitaniae* clones expressing *CauMRR1a* did not differ from that of U04 *mrr1*Δ (4 µg/mL; Table S4), so FLZ MIC alone could not indicate whether this allele is functional in *C. lusitaniae*. One clone expressing each *C. auris MRR1a* allele was chosen at random for the remaining experiments described in this paper: clone #1 for *CauMRR1a^N647T^* and clone #5 for *CauMRR1a*. Using qRT-PCR, we then examined basal expression levels of *C. lusitaniae MGD1* (*CLUG_01281*) and *MDR1* (*CLUG_01938/CLUG_01939*) in the heterologous complements and the U04 *mrr1*Δ parent. Complementation with *CauMRR1a^N647T^* conferred a significant increase in basal expression of both *MGD1* (Fig. 3E) and *MDR1* (Fig. 3F) compared to the *mrr1*Δ parent, while complementation with *CauMRR1a* led to a small, but significant, decrease in expression of both genes relative to *mrr1*Δ (Fig. 3E-F). These results are consistent with our previous observations that *C. lusitaniae* strains expressing certain Mrr1 variants with low basal activity demonstrate lower expression of some Mrr1-regulated genes, including *MDR1* and *MGD1*, compared to an isogenic *mrr1*Δ strain suggesting that Mrr1 has both repressing and activating roles (37, 58). Finally, we assessed the relative MG resistance of the isogenic *C. lusitaniae* strains expressing *CauMRR1a^N647T^* or *CauMRR1a* and the U04 *mrr1*Δ parent. The *CauMRR1a^N647T^* complement grew markedly better in 15 mM MG compared to U04 *mrr1*Δ whereas the *CauMRR1a* complement grew substantially worse than U04 *mrr1*Δ (Fig. 3G), consistent with the pattern of *MGD1* expression we observed in these strains via qRT-PCR. None of the *C. lusitaniae* strains demonstrated growth differences in the YPD control, or in the presence of MG at concentrations of 5 mM or 10 mM (Fig. S4).

### MG induces expression of *MGD1* and *MDR1* in *C. auris* B11221 and AR0390, but not in *C. lusitaniae* strains expressing *C. auris MRR1a* alleles

Next, we compared induction of *MGD1* and *MDR1* by MG in the *C. auris* strains B11221 and AR0390 via qRT-PCR. MG significantly induced expression of *MGD1* by 2.4-fold in *C. auris* strain B11221 and by 4.0-fold in *C. auris* strain AR0390 (Fig. 4A) and expression of *MDR1* by 6.0-fold in B11221 and 9.3-fold in AR0390 (Fig. 4B). AR0390 displayed lower expression of both genes in MG, but a higher fold change compared to B11221, further supporting the hypothesis that the N647T allele is gain-of function.

**Figure 4.**
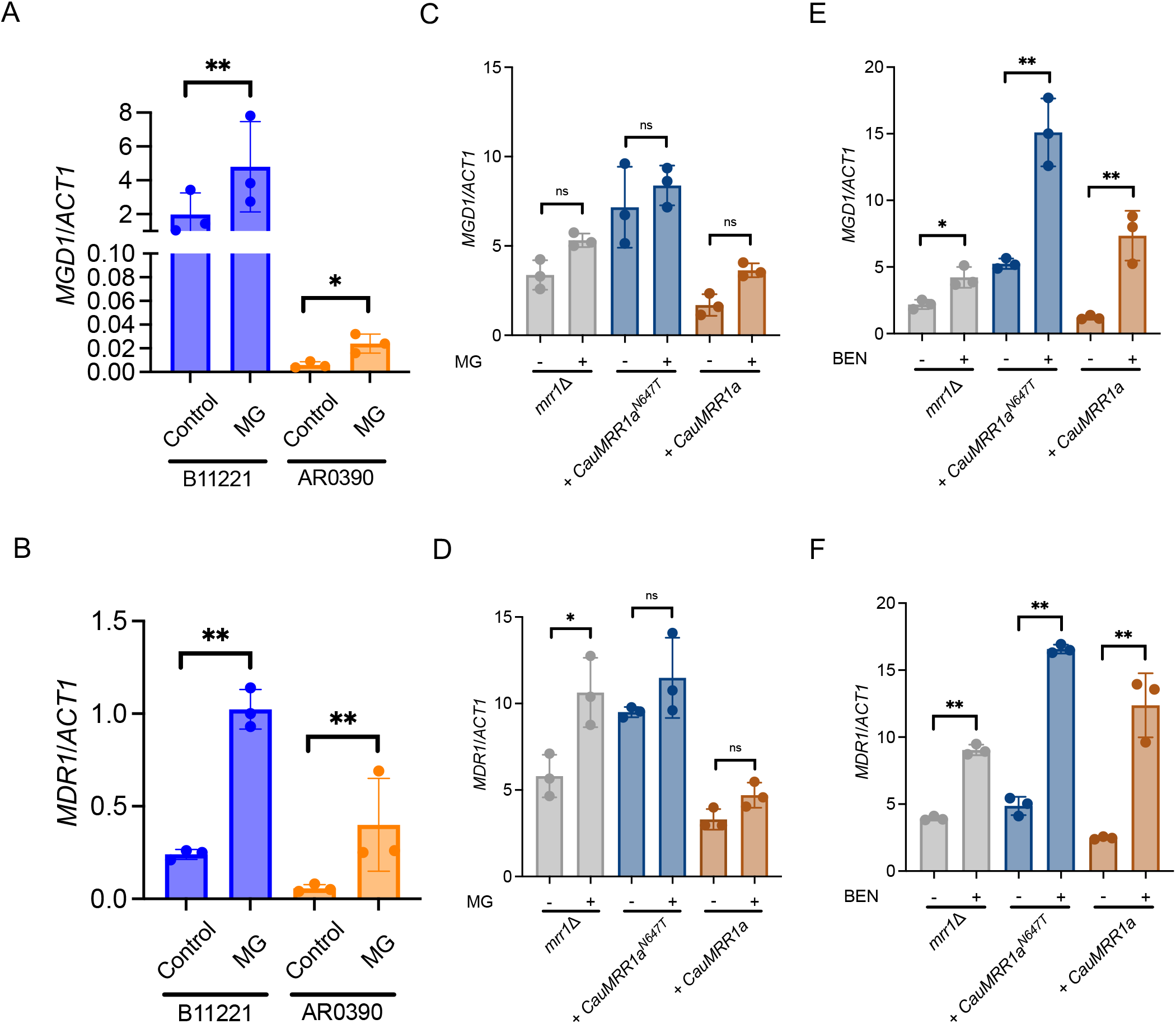
MG induces expression of *MGD1* and *MDR1* in *C. auri*s isolates B11221 and AR0390, but *C. auris MRR1a* is not inducible by MG when heterologously expressed in *C. lusitaniae*. **A-B)** qRT-PCR analysis for expression of *MGD1* **(A)** and *MDR1* **(B)** in exponential-phase cultures of B11221 (blue) or AR0390 (orange) treated with MG as indicated. Data shown represent the mean ± SD for three independent experiments. Ratio paired t-test was used for statistical evaluation; ns p > 0.05, * p < 0.05, ** p < 0.01. **(C-F)** qRT-PCR analysis for expression of *MGD1* **(C,E)** and *MDR1* **(D,F)** in exponential-phase cultures of *C. lusitaniae* U04 *mrr1*Δ (grey) and its derivatives expressing *CauMRR1a^N647T^* (dark blue) or *CauMRR1a* (brown) treated with 5 mM MG for 15 min **(C,D)** or 25 µg/mL BEN for 30 min **(E,F)**. Data shown represent the mean ± SD for three independent experiments. Ratio paired t-test was used for statistical evaluation; ns p > 0.05, * p < 0.05, ** p < 0.01.

Finally, we compared induction of *MGD1* and *MDR1* by MG in the isogenic *C. lusitaniae* strains expressing either *CauMRR1a^N647T^* or *CauMRR1a* and the *mrr1*Δ parent. Additionally, we tested induction by BEN in these strains as a control. While the *mrr1*Δ parent exhibited a significant 1.8-fold induction of *MDR1*, neither *C. lusitaniae* strain expressing a *C. auris* Mrr1a allele demonstrated a significant change in *MGD1* or *MDR1* expression in response to MG (Fig. 4C-D), indicating that *C. auris* Mrr1a may repress *MRR1*-independent MG induction of *MDR1* in *C. lusitaniae* and that induction of *MGD1* by MG in *C. lusitaniae* requires a functional *MRR1* allele from its own species. Treatment with BEN led to significant increase in expression of *MGD1* (Fig. 4E) and *MDR1* (Fig. 4F) in all three *C. lusitaniae* strains. In response to BEN, *MGD1* was induced by 1.9-fold in *mrr1*Δ, 2.9-fold in the *CauMRR1a^N647T^* complement, and 6.1-fold in the *CauMRR1a* complement (Fig. 4E). Likewise, expression of *MDR1* was induced by 2.3-fold in *mrr1*Δ, 3.5-fold in the *CauMRR1a^N647T^* complement, and 5.0-fold in the *CauMRR1a* complement in response to BEN (Fig. 4F). The striking difference in the ability of the *C. lusitaniae* strains expressing *C. auris MRR1a* alleles to respond to BEN versus MG suggests that there are differences in the mechanisms by which BEN and MG induce Mrr1-dependent transcriptional activation and that MG induction of *C. auris* Mrr1a is not supported by *C. lusitaniae* factors. These potential differences are a topic of future study and may shed light on mechanisms of Mrr1 activation in *Candida* species.

### MG and BEN induced Mrr1a-independent transcriptional responses in *C. auris*

We have previously observed heterogeneity in MG resistance as well as MG-induced FLZ resistance among several *C. auris* isolates from different clades (58), and thus we were interested in whether the overall transcriptional response to MG was more similar or different in B11221 and AR0390. AR0390 had greater number of genes differentially expressed by MG compared to B11221; 438 genes were significantly upregulated, and 242 genes were significantly downregulated by MG (see Fig. S5 for the volcano plot of all genes). More genes had a larger fold change in response to MG in AR0390 compared to B11221, including *MGD1* and *MDR1* (Fig. 5A), consistent with the qRT-PCR results in Fig. 4A-B. However, there was a large overlap of 254 genes which were induced by MG in both strains (Fig. 5B), suggesting a common response across these two genetically distinct clades. These commonly induced genes include many with putative roles in amino acid biosynthesis; transmembrane transport; or acquisition and usage of sulfur (Fig. 5C and Table S5). The complete comparison is available in **Supplementary File S6**).

**Figure 5.**
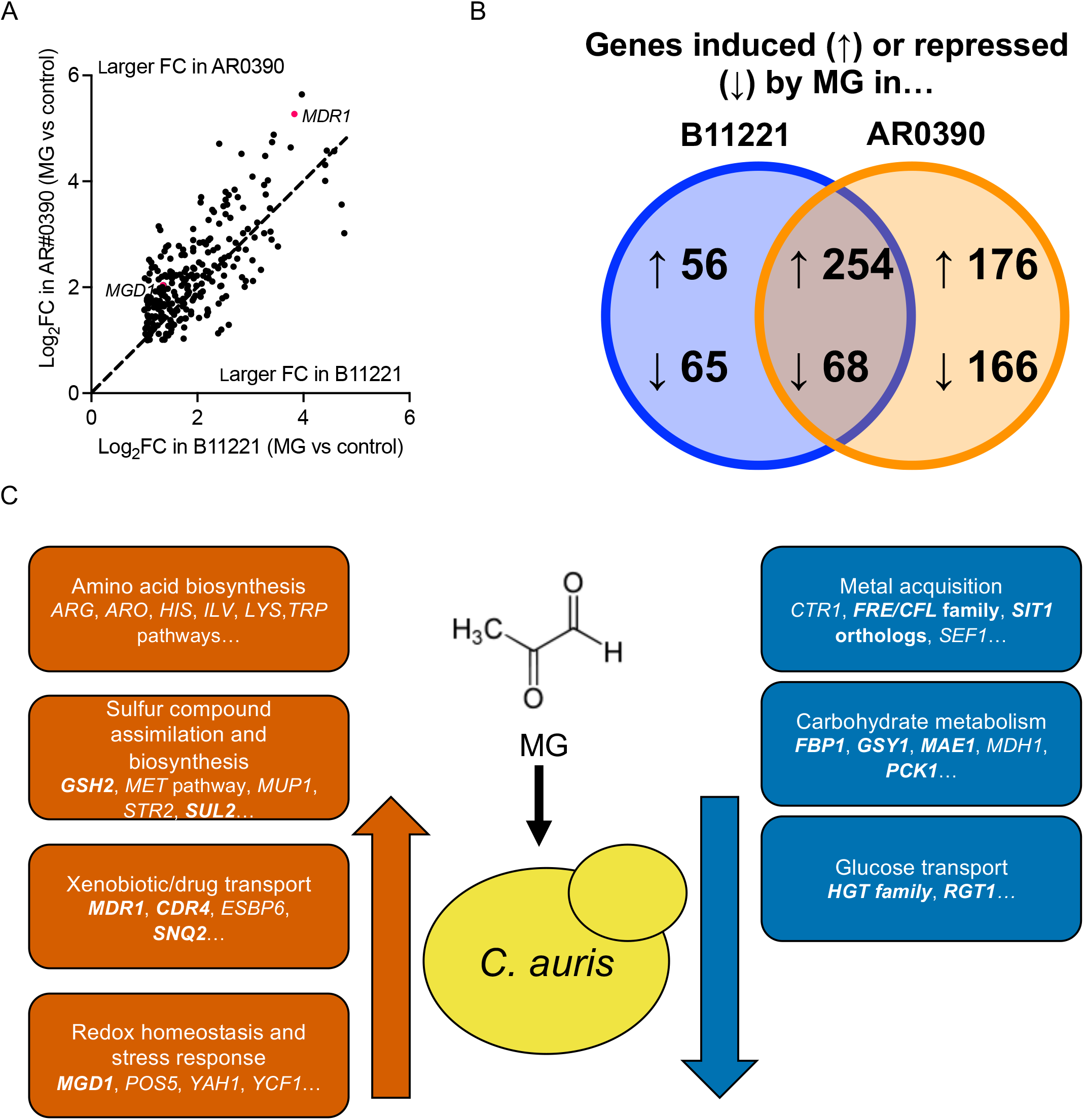
MG induces and represses common pathways across B11221 and AR0390. **A)** Venn diagram of genes with syntenic orthologs between B11221 and AR0390 that were significantly induced (indicated by “up” arrows) or repressed (indicated by “down” arrows) by MG in either or both strains. **B)** Scatter plot of the log_2_FC of genes significantly induced by MG in AR0390 vs the log_2_FC of genes induced by MG in B11221. Only genes with syntenic orthologs between the two strains are shown. Each point represents a single gene; points above the dotted line indicate genes which exhibited a greater Log_2_FC in AR0390, and points below the dotted line indicate genes which exhibited a greater log_2_FC in B11221. *MGD1* and *MDR1* are indicated with red dots for reference. **C)** Graphic summary of major groups of genes that were significantly up- or down-regulated in response to MG in both B11221 and AR0390. Genes in bold text were also up- or down-regulated in response to BEN in B11221.

Only 68 genes with syntenic orthologs across both strains were commonly repressed by MG (Fig 5B). These genes include some with putative roles in metal transport or carbohydrate uptake and metabolism (Fig. 5C and Table S6). We did not observe obvious patterns in genes that were only induced or repressed in one strain, and some genes that are listed as only induced or repressed in one strain were close to the cutoff in the other strain.

The groups of genes that were differentially expressed in response to MG in both B11221 and AR0390 were also evident in the response of B11221 to BEN as well as the response of the *mrr1a*Δ mutant in response to MG and BEN. In B11221, a total of 46 genes exhibited significant induction by both MG and BEN, including *MGD1* and *MDR1*. Many of the 44 other genes have predicted roles in assimilation and biosynthesis of sulfur-containing compounds or xenobiotic transport (Fig. 5C and Table S1). MG also induced expression of many genes with predicted roles in the biosynthesis of amino acids. The two genes most highly upregulated upon MG treatment, in terms of fold change, in this strain were orthologous to the arginine biosynthesis genes *ARG3* (log_2_FC = 4.77) and *ARG1* (log_2_FC = 4.72) (Fig. 2C and Table S1). Conversely, BEN had a limited effect on expression of amino acid biosynthesis genes (Table S1). There were also common themes among the genes that were significantly repressed by both MG and BEN in B11221. Genes that were repressed by both MG and BEN included four orthologs of the *HGT* glucose transporter family, five genes with a predicted role in uptake of iron and/or copper, and *ERG6*, which encodes an enzyme in the ergosterol biosynthesis pathway (Fig. 5C and Table S2). The genes that were repressed by only one stimulus, MG or BEN, also included those involved in ergosterol biosynthesis and the uptake of iron, copper, or glucose (Fig. 5C, Table S2). In general, the transcriptional response of the *mrr1a*Δ mutant to MG and BEN was similar to that of B11221 WT (Fig. S2 and Tables S1-S2). The complete datasets for MG and BEN responses are available in **Supplementary Files S2** and **S3**, respectively.

## Discussion

In this work, we have demonstrated that in *C. auris*, the zinc-cluster transcription factor Mrr1a, which is orthologous to Mrr1 in other *Candida* species, strongly regulates expression of a putative MG reductase *MGD1* in addition to *MDR1*, and that Mrr1a plays a role in MG resistance, highlighting a function of Mrr1 that is distinct from antifungal resistance. We also compared basal global gene expression in B11221 and AR0390 and found that *MDR1*, *MGD1*, and *MGD2* were among the genes significantly more highly expressed in B11221, consistent with the higher MG resistance of this isolate relative to AR0390. These differences were explained by our finding that *MRR1a* from B11221 encoded a higher activity variant than that from AR0390 as evidenced by a higher FLZ MIC, higher expression of *MDR1* and *MGD1*, and higher MG resistance in the strain expressing *CauMRR1a^N647T^* compared to the isogenic strain expressing *CauMRR1a*. The allele from B11221, which contains an N647T amino acid substitution (25, 105) which is in the central region of the regulator where other gain of function substitutions have been found. Both alleles result in induction of *MDR1* and *MGD1* in response to BEN but not to MG in *C*. lusitaniae, suggesting that these two compounds activate Mrr1-dependent transcription through different mechanisms.

Under the conditions tested, Mrr1a regulation in the *C. auris* B11221 background was mainly of *MGD1* and *MDR1*. Homologs of *MDR1* and at least one gene encoding a known or predicted MG reductase are co-regulated by Mrr1 in *C. albicans* (44, 45, 94–96), *C. parapsilosis* (97), and *C. lusitaniae* (37, 40, 58), suggesting that the co-regulation of these two genes has been conserved throughout multiple *Candida* species. Gaining a deeper understanding of the evolutionary and biochemical relationship between methylglyoxal reductases and efflux pumps, particularly Mdr1, may shed light on how *Candida* species sense and respond to environmental or physiological stresses, evade host defense mechanisms, and develop antifungal resistance. In all other *Candida* species with published Mrr1 regulons, however, Mrr1 appears to regulate expression of many more genes than the four we have described here in the *C. auris* strain B11221 (37, 40, 44, 45, 97). The surprisingly small number of *C. auris* genes whose expression was significantly altered by genetic deletion of *MRR1a* may be due to possible redundancy between *MRR1a* and the other two *MRR1* orthologs in *C. auris*, *MRR1b* and *MRR1c*, although further studies would be necessary to test this hypothesis. It is striking, however, that *MRR1a* alone seems to be necessary for expression and induction of *MGD1*, which is further supported by our observation that only the *mrr1a*Δ mutant had a growth defect in MG compared to parental B11221 (Fig. 1A).

Our demonstration of increased basal activity of the *CauMRR1a^N647T^* allele compared to the allele from AR0390 supports the hypothesis put forth by Iyer et al. that the N647T substitution found in many Clade III isolates is a gain-of-function mutation (105). Furthermore, this may explain why deletion of *MRR1a* leads to a mild decrease in azole resistance in B11221, but not in the Clade IV isolate B11243 (46). In *C. albicans*, knocking out gain-of-function *MRR1* causes a significant decrease in FLZ resistance, but knocking out *MRR1* with wild-type transcriptional activity does not alter FLZ resistance (41, 44, 45, 106). Similarly, knocking out gain-of-function *MRR1* in *C. lusitaniae* also decreases FLZ resistance, although knocking out *MRR1* alleles that do not encode a constitutively active protein generally leads to increased FLZ resistance (37).

Although Mrr1a does not appear to play a major role in *C. auris* azole resistance (46), our findings suggest that it contributes to resistance against MG, which may be encountered in the host environment. We have previously shown that Mrr1 also contributes to MG resistance in *C. lusitaniae* in a manner that is partially dependent on *MGD1* and *MGD2* (58). Indeed, gain-of-function mutations in *MRR1* may arise in various *Candida* species due to selective pressures other than azoles. In *C. lusitaniae*, we have reported the emergence of gain-of-function mutations in *MRR1* among isolates from a patient with no prior history of clinical antifungal use (40). In *C. auris*, most sequenced clade III isolates exhibit both the *MRR1a^N647T^* allele and the *ERG11^F126L^* allele (25), the latter of which has been shown to be a major contributor to azole resistance (31). Although it is not known whether the *MRR1a* or *ERG11* mutation occurred first in the clade III lineage, it seems plausible that if the *ERG11* mutation did occur first, evolution of the *MRR1a^N647T^* allele in *C. auris* is likely to be the result of selection for *MGD1* expression and/or an unknown role for Mdr1 that is unrelated to azole resistance. Therefore, we hypothesize that Mrr1 may act, either directly or indirectly, as a response regulator for carbonyl stress in *Candida* species, and future studies will investigate a possible role for Mrr1 in resistance against other physiologically relevant reactive carbonyl compounds.

Curiously, although both variants of *C. auris* Mrr1a were inducible by BEN when expressed in *C. lusitaniae*, they were not inducible by MG under the conditions tested (Fig. 4E-F). One possible hypothesis for this observation is that Mrr1 must interact with at least one particular binding partner to induce transcription in response to MG, and that *C. auris* Mrr1a does not bind efficiently to this *C. lusitaniae* Mrr1-binding protein or complex. Differential requirements for Mrr1-dependent transcriptional activation by chemical stressors have reported in *C. albicans*. For example, the transcription factor Mcm1 is required for Mrr1-dependent induction of *MDR1* in response to BEN but not to H_2_O_2_ (101), and the redox-sensing transcription factor Cap1 is required for *MDR1* induction by H_2_O_2_ and may play a role in *MDR1* induction by BEN (44). Furthermore, gain-of-function Mrr1 in *C. albicans* requires the Swi/Snf chromatin remodeling complex to maintain promoter occupancy, and the kinase Ssn3, which is a subunit of the Mediator complex, may act in opposition to Mrr1 or its coactivators (38). Thus, although *C. auris* Mrr1a can complement Mrr1-dependent basal and BEN-induced expression of *MDR1* and *MGD1* in *C. lusitaniae*, it may be incompatible with certain elements of the *C. lusitaniae* MG-responsive transcriptional machinery. Further studies on the differences between *C. lusitaniae* and *C. auris* Mrr1, particularly in the presence of MG, may elucidate more detailed mechanisms of Mrr1 activation.

In general, we observed substantial upregulation of genes with predicted roles in transmembrane transport, sulfur metabolism, and amino acid biosynthesis in response to MG in all three strains tested. Many genes downregulated in response to MG in all three strains have predicted roles in metal acquisition, particularly iron, and carbohydrate metabolism. In both B11221 WT and *mrr1a*Δ, BEN treatment led to differential expression of similar groups of genes as MG in addition to induction of genes with predicted roles in oxidative stress response. Our studies of the transcriptional response of *C. auris* to MG and BEN contribute to the understanding of how *Candida* species may adapt to oxidative and/or carbonyl stress, two types of stress that a pathogen is likely to encounter in the host environment. In humans, elevated serum MG has been reported in diabetes as well as in renal failure, which are both risk factors for *Candida* infection (107, 108). There is also evidence that neutrophils (109) and macrophages (110, 111) generate MG during the inflammatory response, consistent with elevated levels of MG in sepsis patients (77). In our transcriptomics analysis of three *C. auris* strains exposed to 5 mM MG for 15 min, upregulation of numerous genes involved in amino acid uptake, metabolism, and biosynthesis was one of the most striking responses to MG (Tables S1 for comparison of MG and BEN and **S5** for the comparison of genes induced by MG in B11221 and/or AR0390). In particular, induction of *ARG* genes is interesting considering the report that *C. albicans* upregulates expression of arginine biosynthesis genes when phagocytosed by macrophages or in response to sublethal concentrations of hydrogen peroxide, tert-butyl hydroperoxide, or menadione *in vitro* (112). This induction of *ARG* genes in *C. albicans* by macrophages is dependent on the *gp91^phox^* subunit of the macrophage oxidase, and thus is likely a direct response to oxidative stress rather than arginine depletion (112). In our dataset, *ARG3* and *ARG1* exhibited the highest log_2_FC in response to MG in the B11221 background, independently of *MRR1a* (Table S1). We also observed, in all three *C. auris* strains, induction of several *MET* genes, which are involved in methionine synthesis and are an important branch of sulfur assimilation in yeast. Other genes involved in sulfur acquisition and assimilation that were induced by MG include the sulfate importer *SUL2*, a gene orthologous to both *CYS3* and *STR3* of *S. cerevisiae*, and numerous genes associated with iron-sulfur cluster formation (Table S1). A gene orthologous to *MUP1* of *S. cerevisiae* and *C. albicans* was induced by MG in B11221 WT and AR0390 but fell short of the log_2_FC ≥ 1.00 cutoff in *mrr1a*Δ (Table S1 and **Supplementary File S2**). Induction of genes involved in sulfur metabolism, including the *MET* pathway, *SUL2, CYS3*, *STR3*, and *MUP1*, has previously been observed in *Saccharomyces cerevisiae* exposed to 1g/L acetaldehyde (113), another reactive aldehyde metabolite that is structurally similar to MG. Thus, sulfur acquisition and metabolism may be an important part of the carbonyl stress response in yeast.

In the B11221 background, we observed modest overlap in the genes and groups of genes that were up- or down-regulated in response to either MG or BEN. *MDR1* and *MGD1* were among the genes induced by both compounds, and induction of *MGD1* by either MG or BEN was completely dependent on *MRR1a*. Although BEN, which originated as an agricultural fungicide, is widely recognized as an inducer of expression of Mrr1-regulated genes in *Candida* species (37, 41, 43, 95, 99–104), the mechanism by which this induction occurs is not yet known. BEN is thought to cause oxidative stress in yeast (114, 115), which is consistent with our observation of an upregulation of genes with a predicted role in oxidative stress response in BEN-treated *C. auris* cultures (Table S1). Additionally, in mammalian cells, BEN exposure has been shown to inhibit aldehyde dehydrogenase enzymes (116–119), which may lead to an accumulation of reactive aldehydes, although this possible mechanism has not yet been investigated in fungi.

We also note similarities between the results of our study of MG- and BEN-treated *C. auris* and the recently published transcriptional analysis of the Clade I *C. auris* strain NCPF 8973 exposed to 75 µM farnesol (120). In response to farnesol, the authors reported upregulation of many genes with predicted roles in transmembrane transport, such as *MDR1* and *CDR1*, and downregulation of numerous genes predicted to be involved in metal acquisition and homeostasis, including multiple ferric reductases and iron permeases (120). As farnesol may cause oxidative stress in *Candida* species (120–123) and in *S. cerevisiae* (124, 125), the overlap in transcriptional changes in response to MG, BEN, and farnesol likely provides valuable insight into how *C. auris* and other *Candida* species sense and adapt to physiologically relevant stressors. In fact, MG itself may serve as a stress signal in various organisms. In plants, for example, intracellular MG increases in response to drought (126, 127), salinity (126, 128–131), cold stress (126), heavy metals (128), or phosphorous deficiency (131), and overexpression of certain genes involved in MG detoxification has been shown to enhance salt tolerance in tobacco (126) and in *Brassica juncea* (132). Investigating whether MG detoxification is linked to abiotic stressors such as salt, temperature, or desiccation in *Candida* species would be an interesting avenue of future research, particularly in *C. auris* due to its persistence on hospital surfaces and high salt tolerance.

## Methods

### Strains, media, and growth conditions

The sources of all strains used in this study are listed in Table S7. All strains were stored long term in a final concentration of 25% glycerol at −80°C and freshly streaked onto yeast extract peptone dextrose (YPD) agar (10 g/L yeast extract, 20 g/L peptone, 2% glucose, 1.5% agar) once every seven days and maintained at room temperature. Unless otherwise noted, all overnight cultures were grown in 5 mL YPD liquid medium (10 g/L yeast extract, 20 g/L peptone, 2% glucose) on a rotary wheel at 30°C. Media was supplemented with 25 µg/mL BEN (stock 10 mg/ml in DMSO) or 5 mM, 10 mM, or 15 mM MG (Sigma-Aldrich, 5.55 M) as noted. *E. coli* strains were grown in LB with 15 µg/mL gentamycin (gent).

### Plasmids for complementation of *C. auris MRR1a*

Plasmids for complementing *C. auris MRR1a* into *C. lusitaniae* were created as follows: the open reading frame of *MRR1a* was amplified from the genomic DNA of *C. auris* isolates B11221 (for *CauMRR1a^N647T^*) and AR0390 (for *CauMRR1a*) using a forward primer with homology to the 5’ flank of *C. lusitaniae MRR1* and a reverse primer with homology to the 3’ flank of *C. lusitaniae MRR1* for recombination into the *C. lusitaniae MRR1* complementation plasmid pMQ30*^MRR1-L1191H+Q1197*^* (58). Plasmid *pMQ30^MRR1-L1191H+Q1197*^* was digested with AscI (New England BioLabs) and AgeI-HF (New England BioLabs). The PCR products and digested plasmid were cleaned using the Zymo DNA Clean & Concentrator kit (Zymo Research) and assembled using the *S. cerevisiae* recombination technique described in (133). Recombined plasmids were isolated from *S. cerevisiae* using a yeast plasmid miniprep kit (Zymo Research) before transformation into NEB®5-alpha competent *E. coli* (New England BioLabs). *E. coli* containing pMQ30-derived plasmids were selected for on LB containing 15 µg/mL gentamycin. Plasmids from *E. coli* were isolated using a Zyppy Plasmid Miniprep kit (Zymo Research) and subsequently verified by Sanger sequencing with the Dartmouth College Genomics and Molecular Biology Shared Resources Core. *MRR1a* complementation plasmids containing the correct sequences were linearized with Not1-HF (New England BioLabs), cleaned up with the Zymo DNA Clean & Concentrator kit (Zymo Research) and eluted in molecular biology grade water (Corning) before transformation of 1.5 µg into *C. lusitaniae* strain U04 *mrr1*Δ as described below. All plasmids used and created in this study are listed in Table S7. All primers used in this study are listed in Table S8.

### Transformation of *C. lusitaniae* with *C. auris MRR1a* complementation constructs

Mutants in *C. lusitaniae* were generated using an expression-free CRISPR-Cas9 method as previously described (37, 58, 134). In brief, cells suspended in 1M sorbitol were electroporated immediately following the addition of 1.5 µg of *C. auris MRR1a* complementation plasmid that had been previously linearized with NotI-HF (New England BioLabs) and Cas9 ribonucleoprotein containing crRNA targeting the *NAT1* gene. Transformants were selected on YPD agar containing 600 µg/ml hygromycin B (HygB). Successful transformants were identified via PCR of the *C. lusitaniae MRR1* locus as previously described (37, 58). CRISPR RNAs (crRNAs; IDT) and primers used to validate transformants are listed in Table S8.

### Minimum Inhibitory Concentration (MIC) Assay

MIC assays for FLZ were performed in RPMI-1640 medium (Sigma, containing L-glutamine, 165 mM MOPS, 2% glucose at pH 7) as described in (40) and (58) using the broth microdilution method. The final concentration of FLZ in each well ranged from 64 µg/mL to 0.125 µg/mL. Plates were incubated at 35°C and scored for growth at 24 and 48 hours; the results are reported in Table S4. The MIC was defined as the drug concentration that abolished visible growth compared to a drug-free control.

### Growth Kinetics

Growth kinetic assays were performed as previously described in (58). In brief, exponential-phase cultures of *C. auris* or *C. lusitaniae* were washed and diluted in dH_2_O to an OD_600_ of 1; 60 µL of each diluted cell suspension was added to 5 mL fresh YPD. To each well of a clear 96-well flat-bottom plate (Falcon) was added 100 µL of YPD or YPD with MG at twice the desired final concentration and 100 µL of cell inoculum in YPD. Plates were arranged in technical triplicate for each strain and condition and incubated in a Synergy Neo2 Microplate Reader (BioTek, USA) according to the following protocol: heat to 37°C, start kinetic, read OD_600_ every 60 minutes for 36 hours, end kinetic. Results were calculated in Microsoft Excel and plotted in GraphPad Prism 9.0.0 (GraphPad Software).

### Quantitative Real-Time PCR

Overnight cultures of *C. auris* or *C. lusitaniae* were diluted 1:50 into 5 mL fresh YPD, and grown to for four hours at 37°C. To each culture was added MG to a final concentration of 5 mM (4.5 µL stock), BEN to a final concentration of 25 µg/mL (12.5 µL stock), or 4.5 µL molecular biology grade dH_2_O. Cultures were returned to the roller drum at 37°C for 15 min (MG or dH_2_O) or 30 min (BEN), then centrifuged at 5000 rpm for 5 min. The differences in time of exposure in the experimental scheme was used to maintain consistency with published experiments in other species, and not because of known differences in kinetics of activity for the two inducers. RNA isolation, gDNA removal, cDNA synthesis, and quantitative real-time PCR were performed as previously described (40). Transcripts were normalized to *C. auris* or *C. lusitaniae ACT1* expression as appropriate. Results were calculated in Microsoft Excel and plotted in GraphPad Prism 9.0.0 (GraphPad Software). Primers are listed in Table S8.

### RNA sequencing

Overnight cultures of *C. auris* were diluted to an OD_600_ of 0.1 in 5 mL fresh, pre-warmed YPD, and incubated on a roller drum at 37°C for 5-6 doublings (approx. 6 hours). Cultures were diluted once more to an OD_600_ of 1 in 5 mL fresh, pre-warmed YPD and returned to the roller drum at 37°C for another 5-6 doublings. To each culture was added MG to a final concentration of 5 mM (4.5 µL), BEN to a final concentration of 25 µg/mL (12.5 µL), or 4.5 µL molecular biology grade dH_2_O. Cultures were returned to the roller drum at 37°C for 15 min (MG or dH_2_O) or 30 min (BEN), then centrifuged at 5000 rpm for 5 min. Supernatants were discarded and RNA isolation was performed on cell pellets as described above for qRT-PCR. gDNA was removed from RNA samples as described above. DNA-free RNA samples were sent to the Microbial Genome Sequencing Center (https://www.migscenter.com/) for RNA sequencing.

### Analysis of RNAseq

RNAseq data were analyzed by the Microbial Genome Sequencing Center (https://www.migscenter.com/) as follows: Quality control and adapter trimming was performed with bcl2fastq (https://support.illumina.com/sequencing/sequencing_software/bcl2fastq-conversion-software.html). Read mapping was performed with HISAT2 (135). Read quantification was performed using Subread’s featureCounts (136) functionality. Read counts were loaded into R (https://www.R-project.org/) and normalized using edgeR’s (137) Trimmed Mean of M values (TMM) algorithm. Subsequent values were then converted to counts per million (cpm). Differential expression analysis was performed using edgeR’s Quasi Linear F-Test. In the supplementary files, sheets named “All Quantified Genes” contain the results of the exact test for all genes in addition to the normalized counts per million. Differentially expressed genes were determined using the cutoff of |logFC| > 1 and p < .05.

### Identification of orthologs

Orthologs of *C. auris* genes in *C. albicans*, *C. lusitaniae*, and *S. cerevisiae*, as well as orthologs between B11221 and the Clade I reference strain B8441, were identified using FungiDB (https://fungidb.org) (138, 139).

### Generation of Venn diagrams

Venn diagrams of differentially expressed genes across different strains and conditions were computed using the Venn diagram tool from UGent Bioinformatics & Evolutionary Genomics, which is accessible at https://bioinformatics.psb.ugent.be/webtools/Venn/.

### Statistical Analysis and Figure preparation

All graphs were prepared with GraphPad Prism 9.0.0 (GraphPad Software). Ratio paired t-tests and one-way ANOVA tests were performed in Prism; details on each test are described in the corresponding figure legends. All p-values were two-tailed and p < 0.05 were considered significant for all analyses performed and are indicated with asterisks in the text: * p <0.05, ** p <0.01, *** p <0.001, **** p <0.0001.

### Data availability

The data supporting the findings in this study are available within the paper and its supplemental material and are also available from the corresponding author upon request. The raw sequence reads from the RNA-Seq analysis have been deposited into NCBI sequence read archive under BioProject PRJNA801628.

## Acknowledgements

We thank Joachim Morschhäuser and the FDA-CDC Antimicrobial Resistance Isolate Bank for providing strains. We thank Judith Berman for the pGEM-*URA3* plasmid used for yeast cloning. We thank Elora Demers for primers.

## Author contributions

ARB and DAH conceived and designed the experiments and wrote the paper. ARB performed the experiments. ARB and DAH analyzed the data.

## Funding

This study was supported by grants R01 5R01 AI127548 to DAH. Core services were provided by STANTO19R0 to CFF RDP, P30-DK117469 to DartCF, and P20-GM113132 to BioMT. Sequencing services and specialized equipment were provided by the Genomics and Molecular Biology Shared Resource Core at Dartmouth, NCI Cancer Center Support Grant 5P30 CA023108-41. The content is solely the responsibility of the authors and does not necessarily represent the official views of the NIH.

## Competing interests

The authors have declared that no competing interests exist.

---

**Figure S1.**
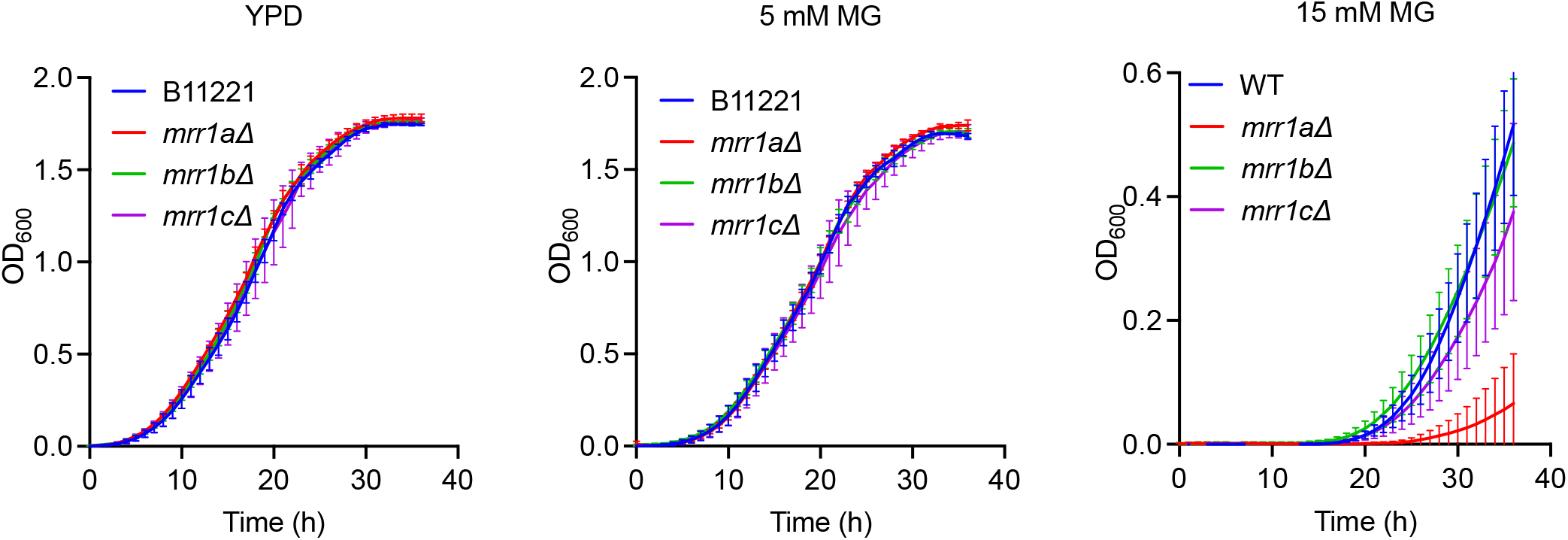
The *mrr1a*Δ mutant has a growth defect in high concentrations of MG, but not at 5 mM MG or in the YPD control. Growth curves of B11221 WT (blue) and its *mrr1a*Δ (red), *mrr1b*Δ (green), and *mrr1c*Δ (purple) derivatives in YPD (left), or YPD supplemented with 5 mM (middle), or 15 mM (right) MG. Data shown represent the mean ± SD for three independent experiments.

**Figure S2.**
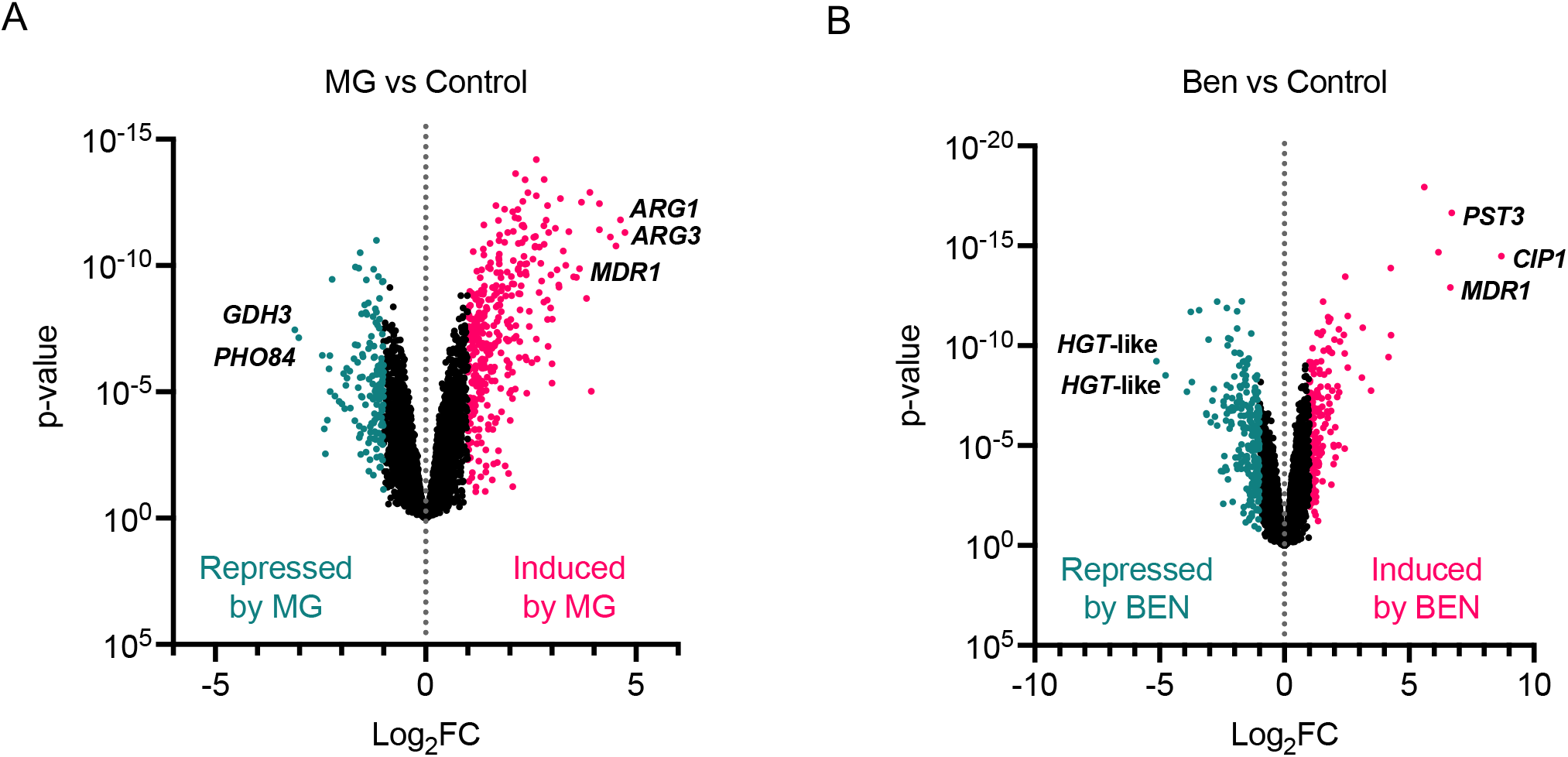
The transcriptional response of *mrr1a*Δ to either MG or BEN is overall similar to that of the B11221 WT parent strain. Volcano plots of all quantified genes in the *mrr1a*Δ mutant treated with either MG **(A)** or BEN **(B)**. Each point represents a single gene; magenta points indicate genes that were significantly upregulated compared to the control condition, teal points indicate genes that were significantly downregulated compared to the control condition. *MDR1* is shown along with the two most up- and down-regulated genes in each condition.

**Figure S3.**
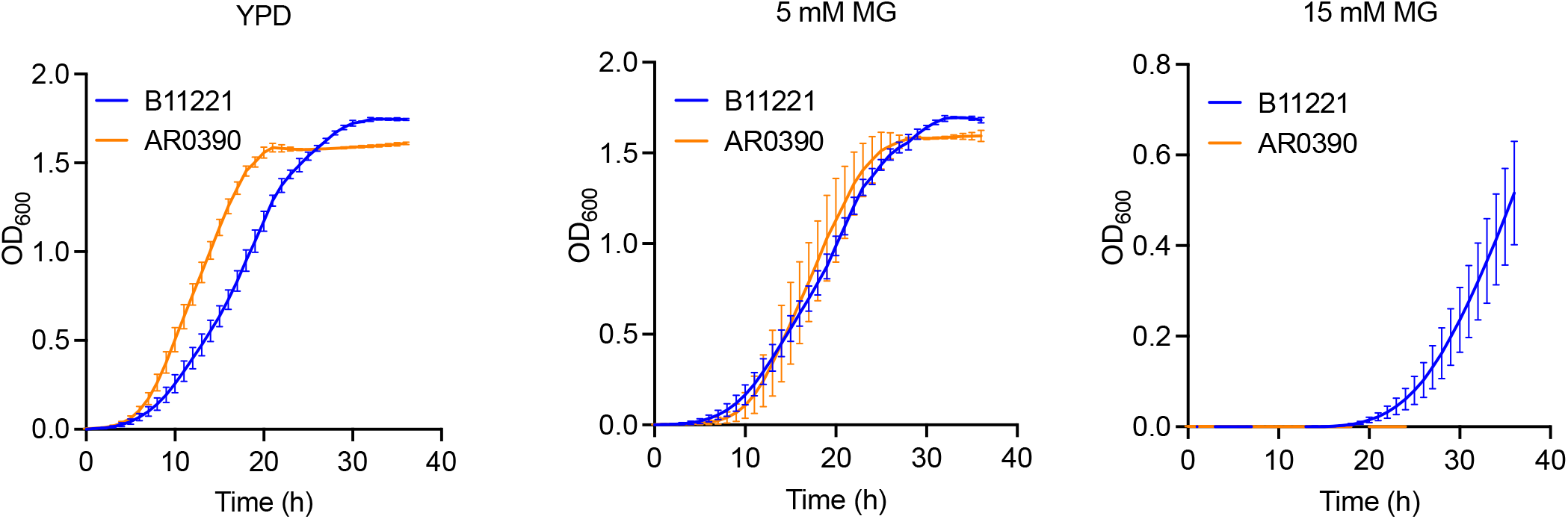
*C. auris* strain AR0390 has a growth advantage over B11221 in YPD but loses that advantage in the presence of increasing concentrations of MG. Growth curves of B11221 (blue) and AR0390 (orange) in YPD (left), or YPD supplemented with 5 mM (middle), or 15 mM (right) MG. Data shown represent the mean ± SD for three independent experiments.

**Figure S4.**
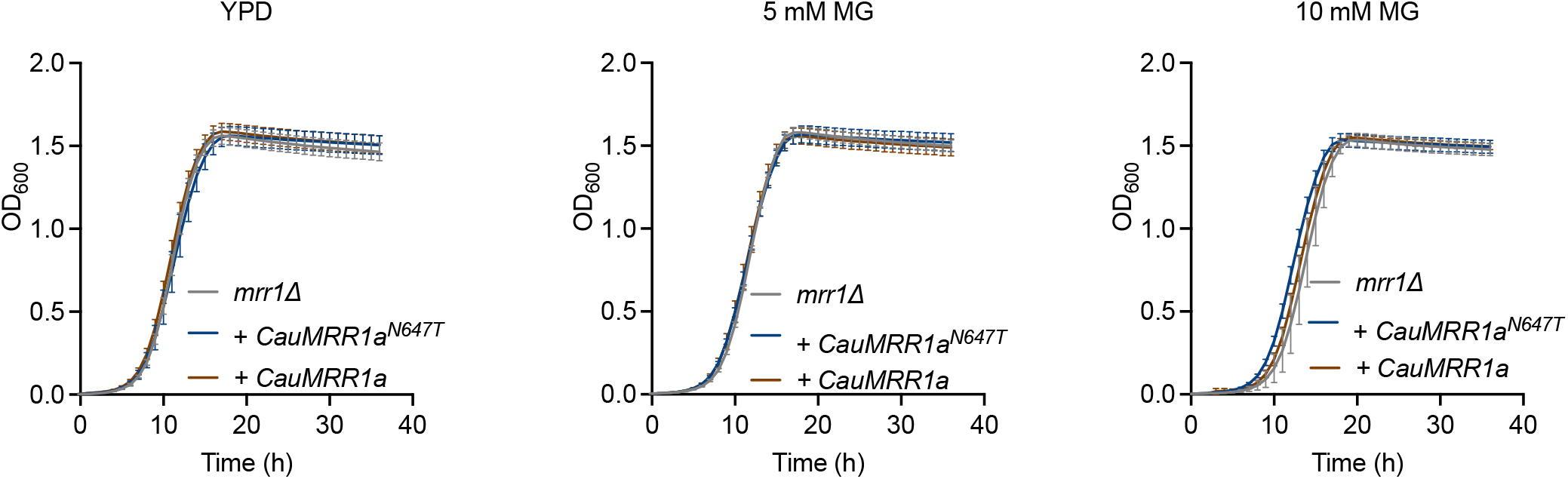
*C. lusitaniae* strains complemented with *CauMRR1aN647T* or *CauMRR1a* do not differ in growth from the mrr1Δ parent at MG concentrations below 15 mM. Growth curves of *C. lusitaniae* U04 *mrr1*Δ (grey) and its derivatives expressing *CauMRR1a^N647T^* (dark blue) or *CauMRR1a* (brown) in YPD (left) or YPD supplemented with 5 mM (middle), or 10 mM (right) MG. Data shown represent the mean ± SD for three independent experiments.

**Figure S5.**
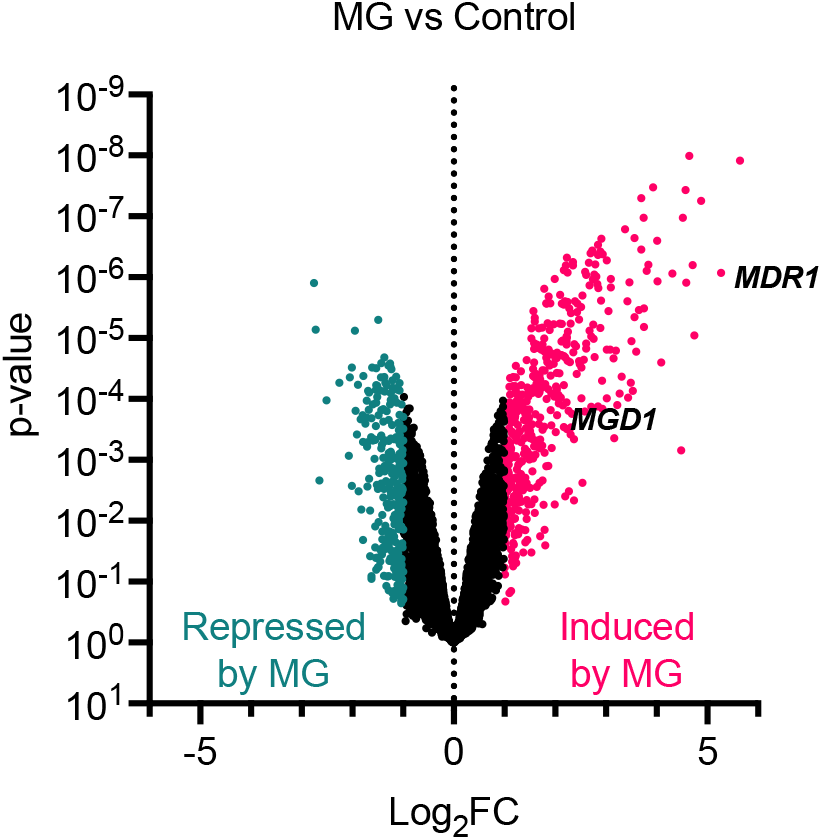
Treatment with 5 mM MG leads to the differential expression of more genes in AR0390 than in B11221. Volcano plot of all quantified genes in AR0390 treated with MG. Each point represents a single gene; magenta points indicate genes that were significantly upregulated compared to the control condition, teal points indicate genes that were significantly downregulated compared to the control condition. *MDR1* and *MGD1* are shown for reference.

**Table S1.**
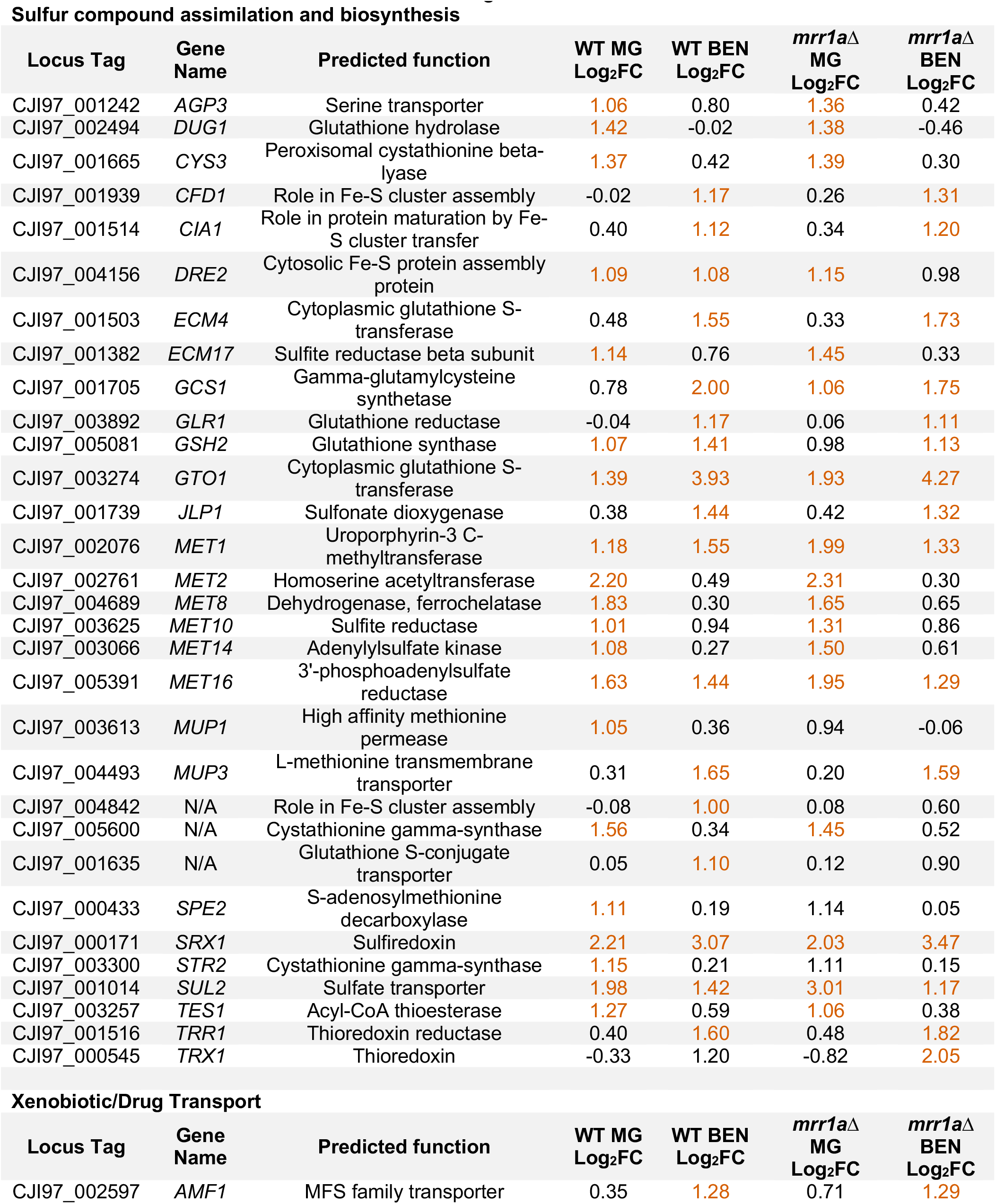
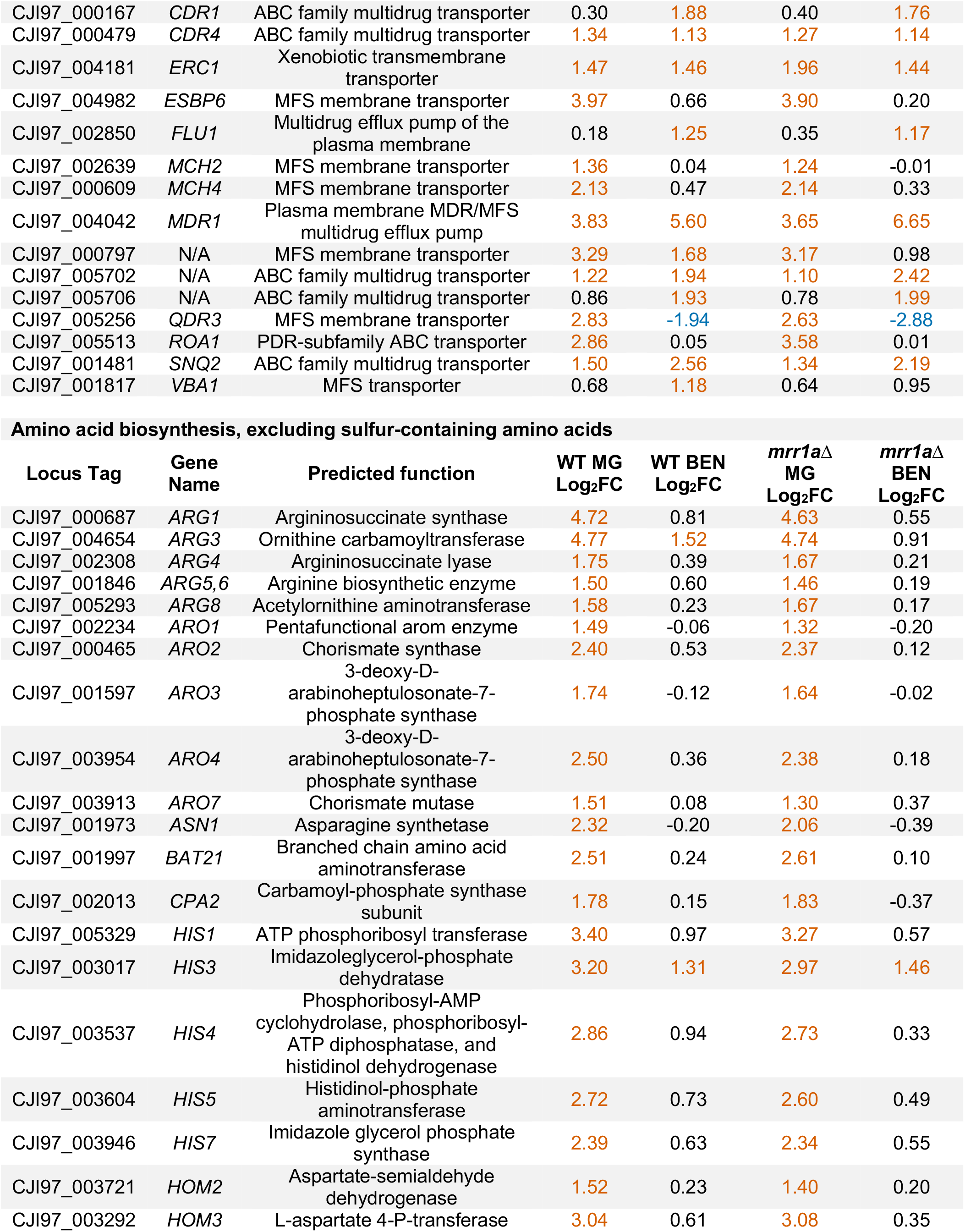
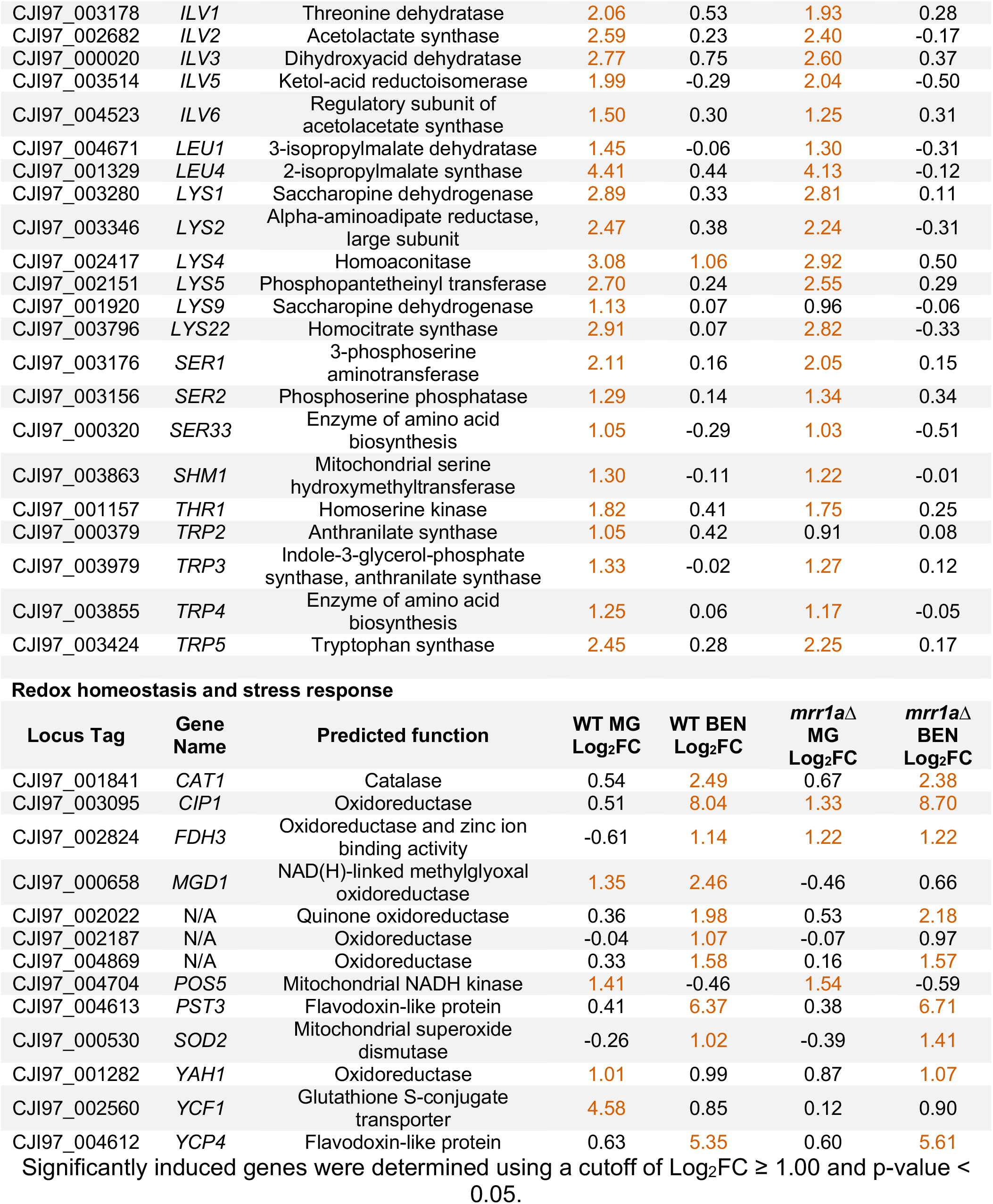
Select genes significantly upregulated by MG and/or BEN in the B11221 background.

**Table S2.**
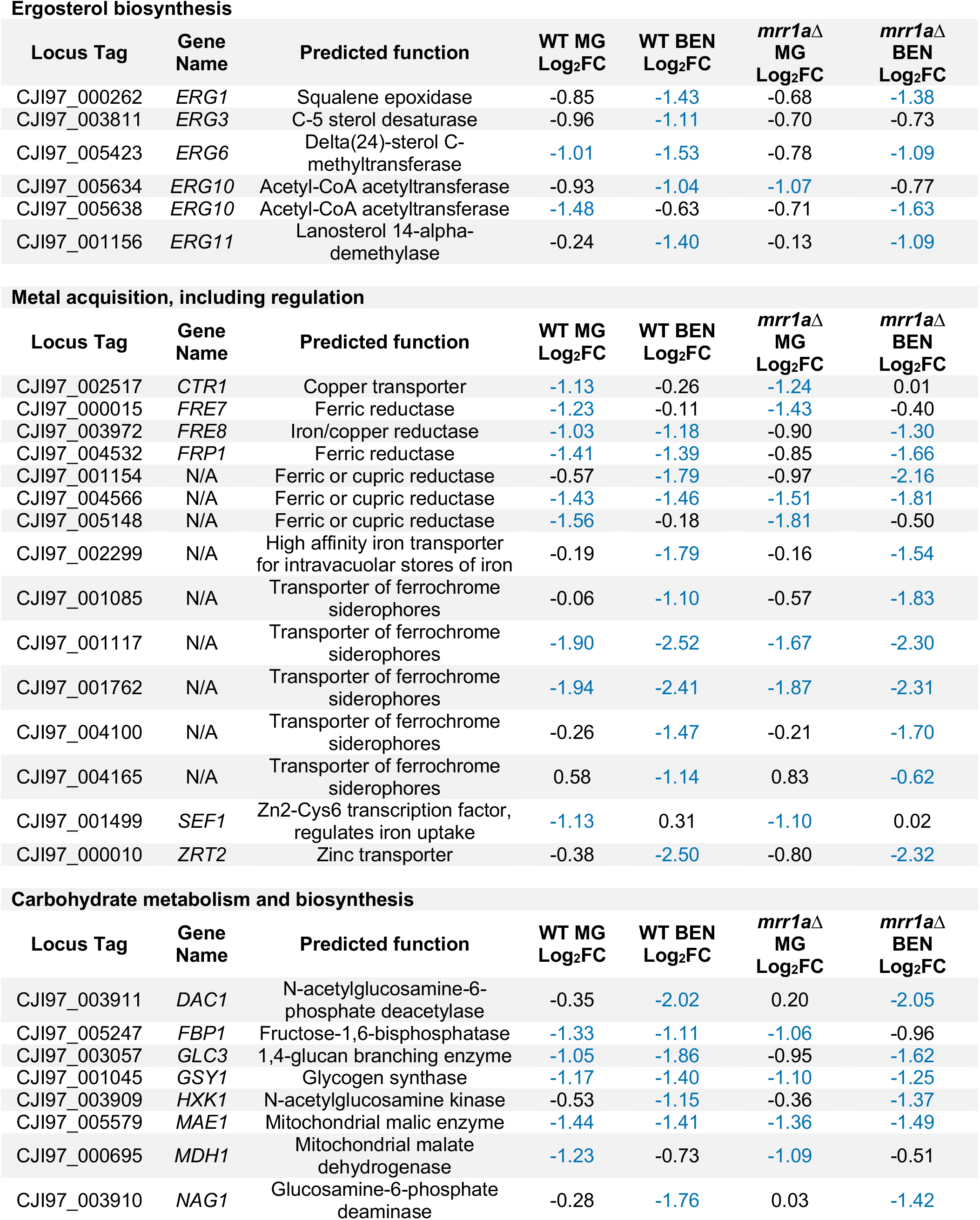
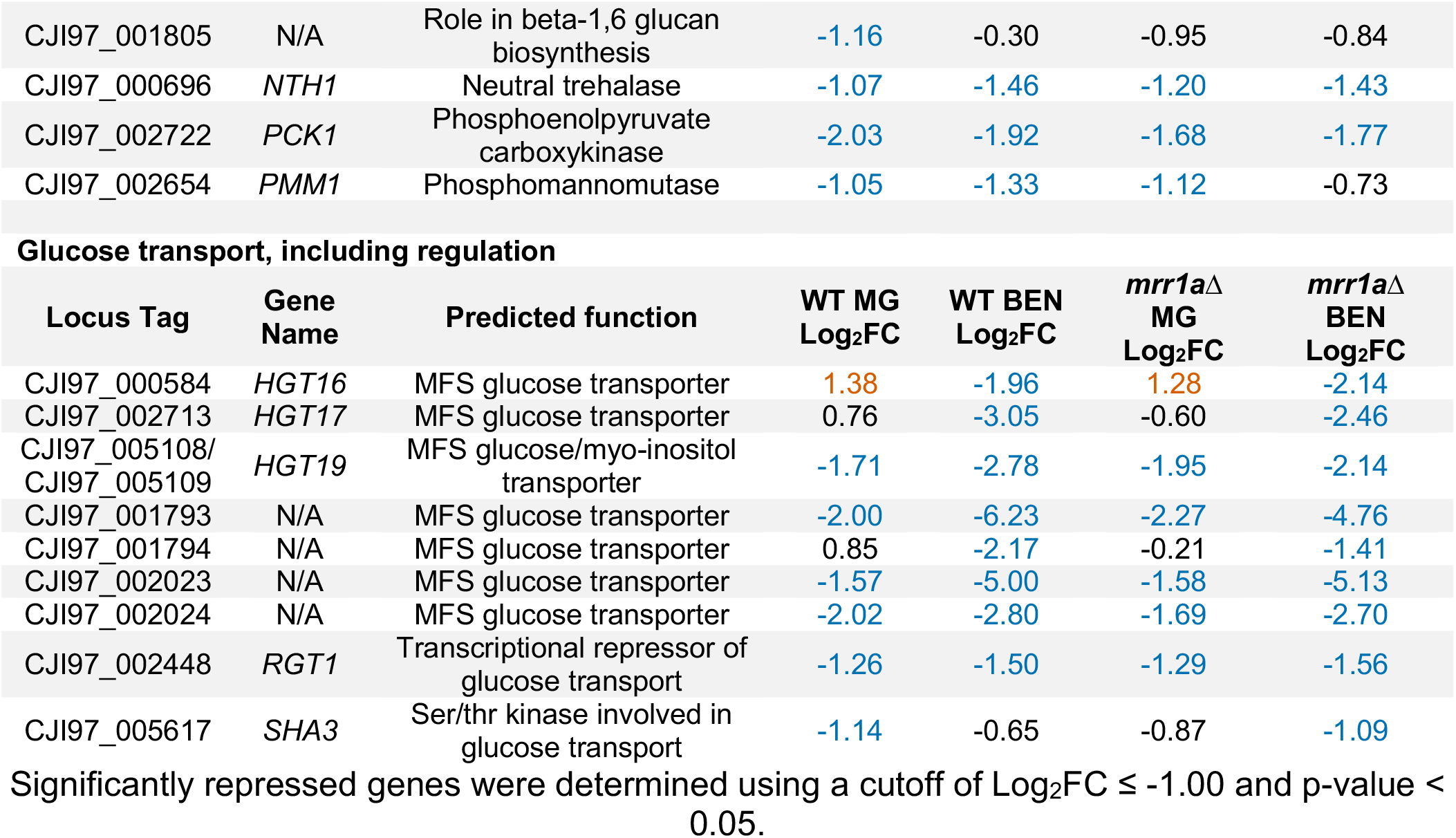
Select genes significantly downregulated by MG and/or BEN in the B11221 background

**Table S3.** Top 20 genes with predicted functions differentially expressed between B11221 and AR0390 in the control condition

**Genes more highly expressed in B11221**
| B11221<br>Locus Tag | AR0390<br>Locus Tag | Gene<br>Name | Predicted Function | Log<sub>2</sub>FC<br>(B11221<br>vs<br>AR0390) | B11221<br>average<br>(CPM) | AR0390<br>average<br>(CPM) |
| --- | --- | --- | --- | --- | --- | --- |
| CJI97_004624 | B9J08_004828 | *MGD2* | NAD(H)-linked methylglyoxal oxidoreductase | 11.29 | 1657 | 0.66 |
| CJI97_000658 | B9J08_000656 | *MGD1* | NAD(H)-linked methylglyoxal oxidoreductase | 8.53 | 4335 | 11.7 |
| CJI97_004768 | B9J08_004684 | N/A | Role in histone deacetylation | 7.87 | 51.9 | 0.22 |
| CJI97_000946 | B9J08_000928 | *AQY1* | Aquaporin water channel, osmotic shock resistance | 7.33 | 581 | 3.60 |
| CJI97_004767 | B9J08_004685 | N/A | Curved DNA-binding protein | 5.36 | 279 | 6.78 |
| CJI97_003833 | B9J08_003761 | N/A | DNA topoisomerase | 5.31 | 96.8 | 2.43 |
| CJI97_002880 | B9J08_002824 | N/A | DNA replication licensing factor required for pre-replication complex assembly | 4.49 | 316 | 14.1 |
| CJI97_004042 | B9J08_003981 | *MDR1* | Plasma membrane MDR/MFS multidrug efflux pump | 4.42 | 190 | 8.88 |
| CJI97_002740 | B9J08_002688 | *FDH1* | Formate dehydrogenase | 4.33 | 131 | 6.50 |
| CJI97_004770 | B9J08_004682 | *ECM42* | Ornithine acetyltransferase | 4.24 | 99.9 | 5.28 |

**Genes more highly expressed in AR0390**
| Locus Tag | AR0390<br>Locus Tag | Gene<br>Name | Predicted Function | Log<sub>2</sub>FC<br>(B11221<br>vs<br>AR0390) | B11221<br>average<br>(CPM) | AR0390<br>average<br>(CPM) |
| --- | --- | --- | --- | --- | --- | --- |
| CJI97_001302 | B9J08_001303 | *BDF1* | Essential chromatin-binding bromodomain protein | -10.21 | 0.48 | 563 |
| CJI97_004515 | B9J08_004451 | N/A | ALS family protein | -9.61 | 1.98 | 1546 |
| CJI97_004556 | B9J08_005565 | *PRD1* | Proteinase | -7.29 | 0.14 | 22.6 |
| CJI97_001865 | B9J08_002322 | *BMH1* | Role in morphology | -7.01 | 12.7 | 1639 |
| CJI97_002974 | B9J08_002900 | *RMD9* | Mitochondrial protein with a predicted role in respiratory growth | -6.86 | 6.41 | 745 |
| CJI97_003073 | B9J08_003002 | N/A | Iron permease | -6.51 | 36.8 | 3350 |
| CJI97_002817 | B9J08_002762 | *INO1* | Inositol-1-phosphate synthase | -5.48 | 28.7 | 1281 |
| CJI97_004514 | B9J08_004450 | *THI13* | Thiamin pyrimidine synthase | -4.95 | 2.34 | 72.4 |
| CJI97_004654 | B9J08_004798 | *ARG3* | Ornithine carbamoyltransferase | -4.95 | 0.70 | 21.5 |
| CJI97_000838 | B9J08_000820 | *SAM4* | S-adenosylmethionine-homocysteine methyltransferase | -4.43 | 1.93 | 41.5 |

Differentially expressed genes were determined using a cutoff of Log<sub>2</sub>FC ≥ 1.00 or ≤ -1.00 and p-value < 0.05.

**Table S4.** Minimum inhibitory concentration (MIC) of fluconazole (FLZ) for *C. lusitaniae* U04 *mrr1*Δ and isogenic strains complemented with either *CauMRR1a^N647T^* or *CauMRR1a* from *C. auris*

| ***C. lusitaniae* strain** | **FLZ MIC (μg/mL)** |  |
| --- | --- | --- |
|  | **24h** | **48h** |
| U04 *mrr1*Δ | 4 | 4 |
| U04 *mrr1*Δ + *CauMRR1a*<sup>N647T</sup> clone # 1 | 16 | 16 |
| U04 *mrr1*Δ + *CauMRR1a*<sup>N647T</sup> clone # 2 | 16 | 16 |
| U04 *mrr1*Δ + *CauMRR1a*<sup>N647T</sup> clone # 8 | 16 | 16 |
| U04 *mrr1*Δ + *CauMRR1a* clone # 4 | 4 | 4 |
| U04 *mrr1*Δ + *CauMRR1a* clone # 5 | 4 | 4 |
| U04 *mrr1*Δ + *CauMRR1a* clone # 7 | 4 | 4 |

**Table S5.** Comparison of select genes induced by MG in B11221 and/or AR0390

| **B11221**<br>**Locus Tag** | **AR0390**<br>**Locus Tag** | **Gene**<br>**Name** | **Predicted Function** | **B11221**<br>**Log<sub>2</sub>FC** | **AR0390**<br>**Log<sub>2</sub>FC** |
| --- | --- | --- | --- | --- | --- |
| CJI97_005163 | B9J08_005078 | *AAH1* | Adenine deaminase, purine salvage and nitrogen catabolism | 0.29 | 1.10 |
| CJI97_002719 | B9J08_002666 | *AGP2* | Amino acid permease | 0.45 | 1.16 |
| CJI97_001242 | B9J08_001362 | *AGP3* | Serine transporter; sulfur assimilation | 1.06 | 0.00 |
| CJI97_004654 | B9J08_004798 | *ARG3* | Ornithine carbamoyltransferase | 4.77 | 3.02 |
| CJI97_003828 | B9J08_003754 | *ARG11* | Ornithine transporter of the mitochondrial inner membrane | 0.94 | 1.70 |
| CJI97_003954 | B9J08_003882 | *ARO4* | 3-deoxy-D-arabinoheptulosonate-7-phosphate synthase; aromatic amino acid biosynthesis | 2.50 | 2.22 |
| CJI97_001997 | B9J08_002453 | *BAT21* | Branched chain amino acid aminotransferase | 2.51 | 2.56 |
| CJI97_005185 | B9J08_005101 | *BNA1* | 3-hydroxyanthranilic acid dioxygenase; NAD biosynthesis | 1.54 | -0.33 |
| CJI97_000479 | B9J08_000479 | *CDR4* | ABC transporter superfamily | 1.34 | 1.01 |
| CJI97_003095 | B9J08_003024 | *CIP1* | Oxidoreductase | 0.51 | 1.30 |
| CJI97_004156 | B9J08_004088 | *DRE2* | Cytosolic Fe-S protein assembly protein | 1.09 | 2.52 |
| CJI97_004181 | B9J08_004118 | *ERC1* | Xenobiotic transmembrane transporter | 1.47 | 0.00 |
| CJI97_002824 | B9J08_002769 | *FDH3* | Oxidoreductase and zinc ion binding activity | 1.25 | 0.90 |
| CJI97_005329 | B9J08_005247 | *HIS1* | ATP phosphoribosyl transferase; histidine biosynthesis | 3.40 | 2.90 |
| CJI97_001933 | B9J08_002388 | *HST6* | ABC transporter related to mammalian P-glycoproteins | 0.66 | 1.18 |
| CJI97_003449 | B9J08_003374 | *ICL1* | Isocitrate lyase; glyoxylate cycle enzyme | 0.79 | 1.33 |
| CJI97_000020 | B9J08_000013 | *ILV3* | Dihydroxyacid dehydratase | 2.77 | 2.73 |
| CJI97_004268 | B9J08_004204 | *JEN1* | Lactate transporter | 1.84 | -0.24 |
| CJI97_001329 | B9J08_001277 | *LEU4* | 2-isopropylmalate synthase | 4.41 | 4.31 |
| CJI97_001920 | B9J08_002375 | *LYS9* | Saccharopine dehydrogenase; lysine biosynthesis | 1.13 | 0.52 |
| CJI97_004689 | B9J08_004763 | *MET8* | Bifunctional dehydrogenase and ferrochelatase; siroheme biosynthesis | 1.83 | 0.94 |
| CJI97_003625 | B9J08_003552 | *MET10* | Sulfite reductase; sulfur amino acid metabolism | 1.01 | 0.90 |
| CJI97_000409 | B9J08_000409 | *MET13* | Methionine biosynthesis protein | 0.04 | 1.65 |
| CJI97_003066 | B9J08_002995 | *MET14* | Adenylylsulfate kinase; sulfur metabolism | 1.08 | 0.93 |
| CJI97_005391 | B9J08_005307 | *MET16* | 3'-phosphoadenylylsulfate reductase; sulfur amino acid metabolism | 1.63 | 1.75 |
| CJI97_004042 | B9J08_003981 | *MDR1* | Plasma membrane MDR/MFS multidrug efflux pump | 3.83 | 5.27 |
| CJI97_000658 | B9J08_000656 | *MGD1* | NAD(H)-linked methylglyoxal oxidoreductase | 1.35 | 2.04 |
| CJI97_004624 | B9J08_004828 | *MGD2* | NAD(H)-linked methylglyoxal oxidoreductase | -0.06 | 1.60 |
| CJI97_002730 | B9J08_002677 | *MIS12* | Mitochondrial C1-tetrahydrofolate synthase precursor | 1.05 | 0.87 |
| CJI97_004904 | B9J08_004548 | *NAR1* | Cytosolic iron-sulfur protein assembly machinery protein | 0.84 | 2.13 |
| CJI97_003488 | B9J08_003413 | *OPT7* | Oligopeptide transporter, may transport GSH or related compounds | 1.09 | 0.90 |
| CJI97_001481 | B9J08_001125 | *SNQ2* | Putative ABC transporter superfamily | 1.50 | 2.25 |
| CJI97_003300 | B9J08_003225 | *STR2* | Cystathionine gamma-synthase; sulfur compound metabolism | 1.15 | 1.32 |
| CJI97_001014 | B9J08_000995 | *SUL2* | Sulfate transporter | 1.98 | 2.03 |
| CJI97_004677 | B9J08_004775 | *TPO3* | Polyamine transporter, MFS-MDR family | 0.91 | 1.24 |
| CJI97_002495 | B9J08_001834 | *TPO4* | Spermidine transporter | 0.48 | 1.49 |
| CJI97_003424 | B9J08_003349 | *TRP5* | Tryptophan synthase | 2.45 | 2.52 |
| CJI97_002560 | B9J08_001899 | *YCF1* | Glutathione S-conjugate transporter | 4.58 | 4.57 |
| CJI97_005451 | B9J08_005368 | *YDJ1* | Type I HSP40 co-chaperone | 0.25 | 1.20 |

Significantly induced genes were determined using a cutoff of $\text{Log}_2\text{FC} \geq 1.00$ and p-value < 0.05.

**Table S6.** Comparison of select genes repressed by MG in B11221 and/or AR0390

| **B11221 Locus Tag** | **AR0390 Locus Tag** | **Gene Name** | **Predicted Function** | **B11221 Log<sub>2</sub>FC** | **AR0390 Log<sub>2</sub>FC** |
| --- | --- | --- | --- | --- | --- |
| CJI97_001435 | B9J08_001171 | *ADH5* | Alcohol dehydrogenase | -1.54 | -1.34 |
| CJI97_002591 | B9J08_001930 | *AOX1* | Alternative oxidase, cyanide-resistant respiration | -1.31 | -0.01 |
| CJI97_004799 | B9J08_004653 | *ARP2* | Component of the Arp2/3 complex | -0.79 | -1.11 |
| CJI97_001469 | B9J08_001137 | *ARP3* | Protein with Myo5p-dependent localization to cortical actin patches at hyphal tip | -0.55 | -1.05 |
| CJI97_000089 | B9J08_000084 | *ATP14* | Mitochondrial F1F0 ATP synthase subunit | -0.45 | -1.02 |
| CJI97_002664 | B9J08_002610 | *ATP17* | Mitochondrial ATPase complex subunit | -0.33 | -1.25 |
| CJI97_003777 | B9J08_003702 | *CDG1* | Cysteine dioxygenases, role in conversion of cysteine to sulfite | -2.12 | -2.65 |
| CJI97_004336 | B9J08_004273 | *COX6* | Cytochrome c oxidase | -0.64 | -1.03 |
| CJI97_002184 | B9J08_001993 | *COX11* | Cytochrome oxidase assembly protein | -0.43 | -1.43 |
| CJI97_004817 | B9J08_004635 | *COX19* | Cytochrome c oxidase assembly protein | -0.39 | -1.16 |
| CJI97_002517 | B9J08_001856 | *CTR1* | Copper transporter | -1.13 | -2.07 |
| CJI97_005423 | B9J08_005340 | *ERG6* | Delta(24)-sterol C-methyltransferase, converts zymosterol to fecosterol, ergosterol biosynthesis | -1.01 | -0.46 |
| CJI97_005321 | B9J08_005239 | *FBA1* | Fructose-bisphosphate aldolase | -0.72 | -1.13 |
| CJI97_005247 | B9J08_005163 | *FBP1* | Fructose-1,6-bisphosphatase, key gluconeogenesis enzyme | -1.33 | -1.48 |
| CJI97_000015 | B9J08_000008 | *FRE7* | Ferric reductase | -1.23 | -0.86 |
| CJI97_003972 | B9J08_004052 | *FRE8* | Iron/copper reductase | -1.03 | -0.23 |
| CJI97_004532 | B9J08_004468 | *FRP1* | Ferric reductase | -1.41 | -0.53 |
| CJI97_002942 | B9J08_002886 | *GIT3* | Glycerophosphocholine permease | -1.18 | -0.13 |
| CJI97_003057 | B9J08_002986 | *GLC3* | 1,4-glucan branching enzyme | -1.05 | -0.93 |
| CJI97_004438 | B9J08_004375 | *GPM1* | Phosphoglycerate mutase | -0.53 | -1.40 |
| CJI97_001045 | B9J08_001025 | *GSY1* | Glycogen synthase | -1.17 | -1.36 |
| CJI97_005108/<br>CJI97_005109 | B9J08_005025 | *HGT19* | MFS glucose/myo-inositol transporter | -1.71 | -0.98 |
| CJI97_002699 | B9J08_002646 | *HXT5* | Sugar transporter | 0.13 | -1.13 |
| CJI97_002817 | B9J08_002762 | *INO1* | Inositol-1-phosphate synthase | -0.36 | -1.41 |
| CJI97_001658 | B9J08_001652 | *JAC1* | ATPase activator activity | -0.25 | -1.04 |
| CJI97_005579 | B9J08_005497 | *MAE1* | Mitochondrial malic enzyme | -1.44 | -1.10 |
| CJI97_000695 | B9J08_000694 | *MDH1* | Mitochondrial malate dehydrogenase | -1.23 | -1.26 |
| CJI97_002683 | B9J08_002630 | *MEP1* | Ammonium permease | -1.81 | -1.38 |
| CJI97_003663 | B9J08_003590 | *MIG2* | Transcription factor involved in glucose repression | -1.17 | -1.44 |
| CJI97_002101 | B9J08_002556 | *MIX14* | Role in aerobic respiration and mitochondrial intermembrane space localization | -0.20 | -1.38 |
| CJI97_002993 | B9J08_002919 | *MLS1* | Malate synthase, glyoxylate cycle enzyme | -0.84 | -1.11 |
| CJI97_001141 | B9J08_001463 | *MYO1* | Component of actomyosin ring at neck of newly emerged bud | -0.98 | -1.22 |
| CJI97_001117 | B9J08_001487 | N/A | Transporter of ferrochrome siderophores | -1.90 | -1.06 |
| CJI97_001762 | B9J08_001547 | N/A | Transporter of ferrochrome siderophores | -1.94 | -1.07 |
| CJI97_000596 | B9J08_000675 | N/A | Adhesin-like protein | -1.12 | 0.15 |
| CJI97_002126 | B9J08_002582 | N/A | Adhesin-like protein | -1.48 | 1.00 |
| CJI97_003987 | B9J08_004037 | N/A | Adhesin-like protein | -1.19 | -0.35 |
| CJI97_004240 | B9J08_004176 | N/A | Secreted lipase | -1.69 | -0.78 |
| CJI97_001776 | B9J08_001533 | N/A | NAD-aldehyde dehydrogenase | -1.57 | -1.55 |
| CJI97_003161 | B9J08_003088 | N/A | NAD-aldehyde dehydrogenase | -1.76 | -1.89 |
| CJI97_001793 | B9J08_002250 | N/A | MFS glucose transporter | -2.00 | -1.41 |
| CJI97_002024 | B9J08_002481 | N/A | MFS glucose transporter | -2.02 | -1.52 |
| CJI97_004566 | B9J08_004886 | N/A | Protein similar to ferric reductases and cupric reductases | -1.43 | -1.43 |
| CJI97_005148 | B9J08_005064 | N/A | Protein similar to ferric reductases and cupric reductases | -1.56 | -1.84 |
| CJI97_000696 | B9J08_000695 | *NTH1* | Neutral trehalase | -1.07 | -0.98 |
| CJI97_002722 | B9J08_002669 | *PCK1* | Phosphoenolpyruvate carboxykinase | -2.03 | -2.01 |
| CJI97_002521 | B9J08_001860 | *PGK1* | Phosphoglycerate kinase | -0.77 | -1.19 |
| CJI97_001140 | B9J08_001464 | *PHO84* | High-affinity phosphate transporter | -2.74 | -2.72 |
| CJI97_001580 | B9J08_002202 | *PHO89* | Phosphate permease | -1.48 | -0.73 |
| CJI97_001697 | B9J08_001613 | *PHO100* | Putative inducible acid phosphatase | -1.38 | -1.60 |
| CJI97_004666 | B9J08_004786 | *PIR1* | 1,3-beta-glucan-linked cell wall protein | -0.44 | -1.12 |
| CJI97_002654 | B9J08_002600 | *PMM1* | Phosphomannomutase, enzyme of O- and N-linked mannosylation | -1.05 | -0.94 |
| CJI97_002321 | B9J08_002130 | *PUT1* | Putative proline oxidase | -2.49 | -1.79 |
| CJI97_004379 | B9J08_004317 | *PUT2* | Putative delta-1-pyrroline-5-carboxylate dehydrogenase | -1.01 | -1.45 |
| CJI97_002448 | B9J08_001787 | *RGT1* | Transcriptional repressor involved in the regulation of glucose transporter genes | -1.26 | -1.62 |
| CJI97_002974 | B9J08_002900 | *RMD9* | Mitochondrial protein with a predicted role in respiratory growth | -0.25 | -1.09 |
| CJI97_004415 | B9J08_004352 | *SAH1* | S-adenosyl-L-homocysteine hydrolase | -1.46 | -1.22 |
| CJI97_004940 | B9J08_004512 | *SCO1* | Copper transporter | -0.79 | -1.06 |
| CJI97_001499 | B9J08_001107 | *SEF1* | Zn2-Cys6 transcription factor, regulates iron uptake | -1.13 | -0.81 |
| CJI97_005617 | B9J08_005567 | *SHA3* | Ser/thr kinase involved in glucose transport | -1.14 | 0.10 |
| CJI97_002536 | B9J08_001875 | *TPI1* | Triose-phosphate isomerase | -0.36 | -1.42 |
| CJI97_003198 | B9J08_003126 | *QCR8* | Ubiquinol cytochrome c reductase | -0.22 | -1.12 |
| CJI97_002481 | B9J08_001820 | *QCR10* | Ubiquinol-cytochrome-c reductase | -0.31 | -1.01 |
| CJI97_003997 | B9J08_004027 | *WOR1* | Transcription factor of white-opaque phenotypic switching in *C. albicans* | -1.22 | -0.33 |

Significantly repressed genes were determined using a cutoff of $\text{Log}_2\text{FC} \leq -1.00$ and $p\text{-value} < 0.05$ .

**Table S7.** Strains used in this study

| Strain | Lab # | Species | Parent | Relevant Characteristics or Genotype | Source |
| --- | --- | --- | --- | --- | --- |
| **Fungal Strains** |  |  |  |  |  |
| AR0390 | DH2777 | *C. auris* |  | Clinical isolate, clade I | (1) |
| B11221 | DH3880 | *C. auris* |  | Clinical isolate, clade III | (2) |
| *mrr1a*Δ | DH3881 | *C. auris* | B11221 | *mrr1a*Δ::caSAT1 | (2) |
| *mrr1b*Δ | DH3882 | *C. auris* | B11221 | *mrr1b*Δ::caSAT1 | (2) |
| *mrr1c*Δ | DH3883 | *C. auris* | B11221 | *mrr1c*Δ::caSAT1 | (2) |
| U04 *mrr1*Δ | DH3306 | *C. lusitaniae* | U04 | *mrr1*Δ::NAT1 | (3) |
| U04 *mrr1*Δ +<br>*CauMRR1a*<sup>N647T</sup> clone #1 | DH3914 | *C. lusitaniae* | U04 *mrr1*Δ | *CauMRR1a*<sup>N647T</sup>-HygB | This study |
| U04 *mrr1*Δ +<br>*CauMRR1a*<sup>N647T</sup> clone #2 | DH3915 | *C. lusitaniae* | U04 *mrr1*Δ | *CauMRR1a*<sup>N647T</sup>-HygB | This study |
| U04 *mrr1*Δ +<br>*CauMRR1a*<sup>N647T</sup> clone #8 | DH3916 | *C. lusitaniae* | U04 *mrr1*Δ | *CauMRR1a*<sup>N647T</sup>-HygB | This study |
| U04 *mrr1*Δ + *CauMRR1a*<br>clone #4 | DH3917 | *C. lusitaniae* | U04 *mrr1*Δ | *CauMRR1a*-HygB | This study |
| U04 *mrr1*Δ + *CauMRR1a*<br>clone #5 | DH3918 | *C. lusitaniae* | U04 *mrr1*Δ | *CauMRR1a*-HygB | This study |
| U04 *mrr1*Δ + *CauMRR1a*<br>clone #7 | DH3919 | *C. lusitaniae* | U04 *mrr1*Δ | *CauMRR1a*-HygB | This study |
| **Plasmids in *E. coli* (DH5α)** |  |  |  |  |  |
| pMQ30<sup>MRR1-L1191H+Q1197*</sup> | DH3829 | *E. coli* |  | *MRR1*<sup>L1191H+Q1197*</sup>-HygB complementation, Gent<sup>R</sup> | (5) |
| pMQ30<sup>CauMRR1a<sup>N647T</sup></sup> | DH3912 | *E. coli* |  | *CauMRR1a*<sup>N647T</sup>-HygB complementation, Gent<sup>R</sup> | This study |
| pMQ30<sup>CauMRR1a</sup> | DH3913 | *E. coli* |  | *CauMRR1a*-HygB complementation, Gent<sup>R</sup> | This study |

**Table S8.**
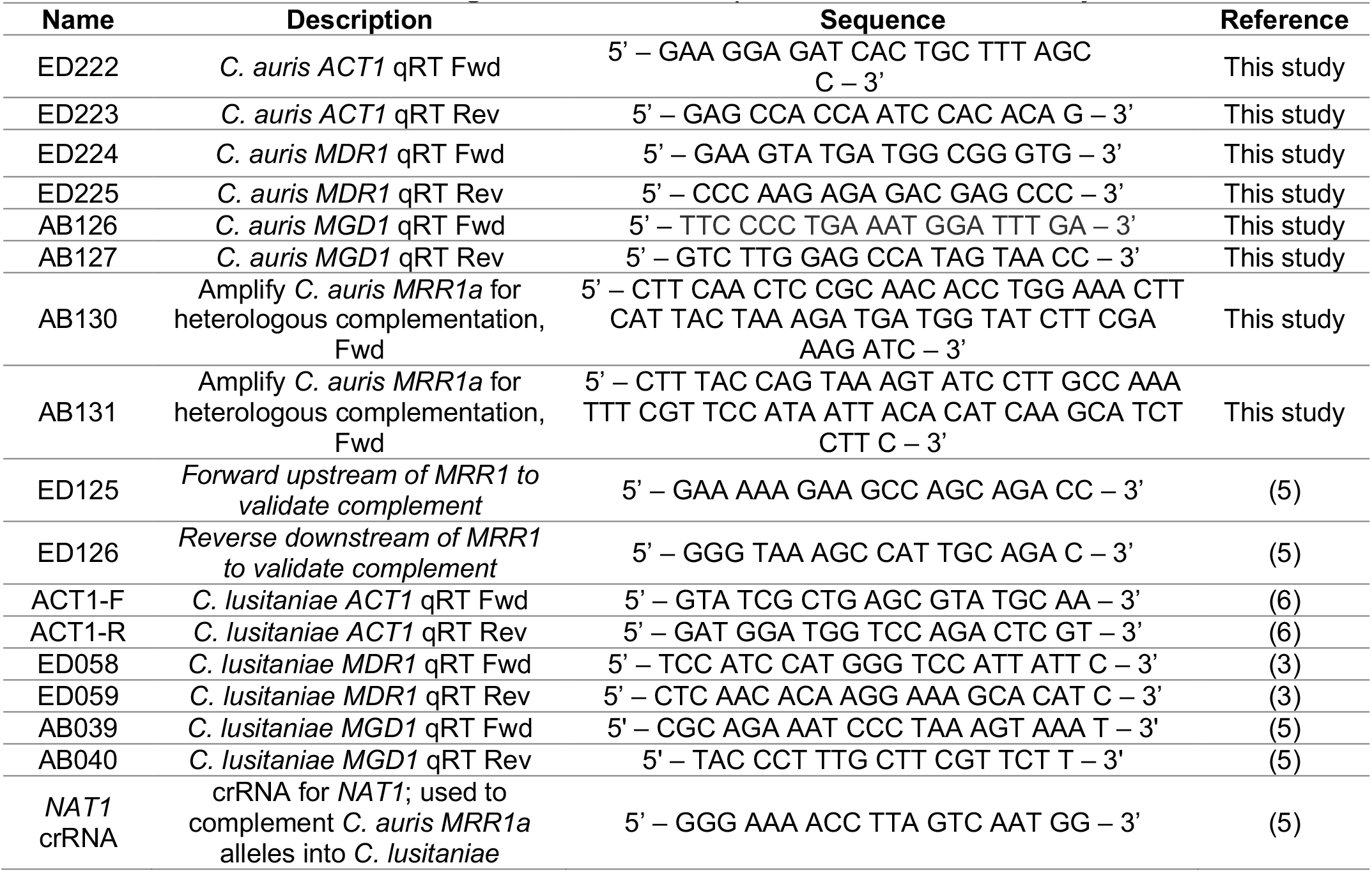
Oligonucleotides and primers used in this study

